# PLCγ2 deficiency compromises systemic immune tolerance and erodes myelin homeostasis while enhancing oxidative metabolism in the mouse brain

**DOI:** 10.64898/2026.07.13.738356

**Authors:** Eduardo Gutierrez-Kuri, Juliet Lynn Marie Garcia-Rogers, Jaid Perez, Sabrina Smith, Matthew R. Kenwood, Kahealani S. Archuleta, Yangming Xiao, Gabriela Campos, Savannah Barannikov, Hu Wang, Sammy Pardo, Trevor B. Romsdahl, Henry Miller, Ann M. Stowe, Russell William, Mark Goldberg, Xianlin Han, Kevin F. Bieniek, Susan T. Weintraub, Ann V. Griffith, Sarah C. Hopp, Juan Pablo Palavicini

## Abstract

**Background:** Phospholipase C gamma-2 (PLCγ2) catalyzes the hydrolysis of the membrane phosphatidylinositol-4,5-bisphosphate (PIP_2_) to form diacylglycerol (DAG) and inositol trisphosphate (IP_3_), feeding into diverse downstream signaling pathways. *PLCG2* polymorphisms have been associated with reduced and/or increased risk of Alzheimer’s disease (AD) and related dementias, longevity, autoinflammation, and immune disorders. In the brain, PLCγ2 is expressed in microglia, and other neuroimmune and vascular interface populations, yet its role in brain homeostasis remains incompletely defined.

**Methods:** We analyzed the brains of three-month-old *Plcg2* wild-type (WT), heterozygous (Het KO) and homozygous knockout (Homo KO) littermate mice modeling human PLCG2 loss-of-function risk alleles linked to AD risk using a multiomic approach that included lipidomics, metabolomics, proteomics, and transcriptomics, together with immunofluorescence, as well as flow-cytometric profiling of peripheral and brain-draining immune compartments.

**Results:** *Plcg2* deficiency substantially impaired early survival and produced splenomegaly without increasing total spleen cellularity, instead shifting spleen composition toward myeloid/innate-enriched cells and away from B cells, with expansion of age-associated B-cell (ABC-like) subsets and parallel reductions in CD4 and CD8 regulatory T cells in spleen and cervical lymph nodes. Brain lipidomics revealed selective depletion of PIP_2_, despite very low bulk PLCγ2 protein abundance relative to other PLC family members. PLCγ2 loss led to significant reductions in myelin-enriched lipid classes and myelin/paranode-associated proteins, accompanied by compensatory upregulation of oligodendrocyte/myelin genes, and modest shifts in microglial, lysosomal, complement, and oxidative metabolism pathways by NanoString and DIA-MS. Targeted acylcarnitine profiling demonstrated reprogramming of brain oxidative metabolism, with increased short-, medium-, and long-chain acylcarnitines and enrichment of mitochondrial matrix fatty-acid and amino-acid catabolic enzymes in Homo KO brains.

**Conclusions:** Loss of PLCγ2 installs a coordinated program that compromises systemic immune tolerance and subtly erodes central myelin and phosphoinositide homeostasis while enhancing brain oxidative metabolism, effects that extend beyond microglial phagocytic signaling and may underlie increased vulnerability to AD pathology and aging, providing a mechanistic framework for how *PLCG2* variation may link systemic immune regulation, white-matter integrity, and neurodegenerative risk.

**Limitations:** Because constitutive *Plcg2* Homo KO mice display high early mortality and intestinal vascular abnormalities, observed phenotypes may reflect developmental compensation and may not fully recapitulate protective human *PLCG2* variants.

## Background

Alzheimer’s disease (AD), the most common cause of dementia, is an age-related neurodegenerative disorder characterized by progressive memory loss and a decline of cognitive function. Histopathological hallmarks of AD include the accumulation of extracellular amyloid plaques and intracellular neurofibrillary tau tangles in the brain. While recent genome-wide association studies have identified several novel genetic variants associated with enhanced or reduced risk for developing AD **[1–3]**, pointing towards microglia/immune-related and lipid metabolism genes, we still need to understand how these loci modulate dementia symptoms.

Multiple *PLCG2* gene polymorphisms have been recently associated with AD. These include *PLCG2* P522R rare variant (rs72824905) that provides robust protection against AD (odds ratio ≈ 0.68 **[1, 4]**, comparable to having one APOE2 allele **[5]**; the very rare *PLCG2* Q816X and R953X loss-of-function (LoF) variants that reduce *PLCG2* expression by ∼50% and confer substantial risk for AD (OR ≈ 10 **[6]**, comparable to having two APOE4 alleles); and other rare PLCG2 variants, such as M28L (rs61749044) that confers a modest increased risk of AD (OR ≈ 1.16) **[1, 4, 7–13]**. Thus, *PLCG2* belongs to a rare class of AD genes with both protective and risk variants, previously exemplified mainly by *APP* and *APOE*, yet remains less well characterized and appreciated than these canonical loci despite its strong genetic support. In addition, *PLCG2* expression in the brain has been associated with both amyloid and tau pathology **[8, 14]**. The phospholipase C-γ-2 (PLCγ2) P522R protective variant is a functional hypermorph **[15–19]**, while M28L is predicted to be a functional hypomorph **[20]**. Expression of PLCγ2 M28L in the mouse brain led to a transcriptomic profile that correlated with human AD modules and led to increased microglia density when mice were fed a high-fat diet **[20]**. Other more strongly hypermorphic *PLCG2* variants are associated with autoimmune PLCγ2-associated antibody deficiency and immune dysregulation (APLAID) **[18]**, while LoF *PLCG2* variants are also associated with common variable immunodeficiency (CVID), PLAID, and familial cold autoinflammatory syndrome (FCAS3) **[21]**. However, the consequences of PLCγ2 loss-of-function in the brain remain largely unknown.

PLCγ2 is primarily expressed in lymphoid tissue (bone marrow, thymus, spleen and lymph nodes), where it functions immediately downstream of the B-cell receptor (BCR) and other ITAM-coupled receptors [22–24]. Genetic or acquired defects in PLCγ2 signaling lead to breakdown of peripheral immune tolerance, autoimmunity, and altered responses to infection in humans and mice. Age-associated B cells (ABCs; CD11c⁺T-bet⁺) and regulatory T cells (Tregs) are key regulators of immune tolerance in aging, and their expansion or loss, respectively, has been implicated in age-related autoimmunity and chronic inflammation [25–29]. These observations raise the possibility that PLCγ2 LoF variants linked to AD risk and longevity may influence disease trajectories in part through systemic immune tolerance mechanisms, including in lymphoid tissues that drain the central nervous system.

*PLCG2* gene expression in the mammalian brain is very low under physiological conditions, yet it is induced in AD brains and AD-like animal models [12, 30]. Within the brain, PLCγ2 is primarily but not exclusively expressed in microglia, where it serves as an important signaling node downstream of TREM2 and CD33 **[31, 32]**, both also AD risk genes **[33–35]**. Public single-cell RNA-seq atlases of the mouse brain indicate that *Plcg2* expression is concentrated in microglia and other neuroimmune and vascular interface populations, consistent with a role for PLCγ2 in coordinating signals at brain–immune and brain–vascular borders. However, it is not known whether this increased *PLCG2* expression in AD reflects enhanced PLCγ2 enzymatic activity or instead compensatory or maladaptive changes in a low-abundance signaling node. Given the localization of PLCγ2 to neuroimmune and vascular interfaces, brain-derived signals are well positioned to influence, and be influenced by, immune circuits in draining lymphoid tissues such as cervical lymph nodes.

PLCγ2 catalyzes the hydrolysis phosphatidylinositol-4,5-bisphosphate (PIP_2_) to diacylglycerol (DAG) and inositol trisphosphate (IP_3_). DAG and IP_3_ feed into downstream signaling pathways involving protein kinase C (PKC), lipid biosynthesis, and intracellular calcium regulation. The TREM2-PLCγ2 signaling axis has been proposed to regulate phagocytosis, secretion of cytokines and chemokines, cell survival and proliferation in microglia **[17, 36, 37]**. Beyond microglia, phosphoinositides including PIP_2_ are abundant in CNS myelin and have been implicated in myelin protein trafficking, cytoskeletal coupling, and paranodal organization [38–41], and multiple myelin-enriched lipid classes (such as galactosylceramides, sulfatides, and plasmalogens) are increasingly recognized as vulnerable nodes in AD [42, 43].

Peroxisomal and mitochondrial oxidative metabolism, including fatty acid β-oxidation, branched-chain fatty acid and amino acid oxidation, and associated carboxylic acid pathways, play an essential role in myelinated tissues and white matter maintenance, and disturbances in these pathways have been linked to myelin pathology, ferroptotic vulnerability, and aging-related brain changes. Moreover, alterations in acylcarnitine profiles and oxidative metabolism have been observed in aging and AD brains, suggesting that shifts in oxidative fuel handling may accompany or compensate for lipid and myelin perturbations.

In the studies described herein, we tested the hypothesis that haploinsufficiency or full deficiency of *Plcg2* in mice, as a model for the novel human *PLCG2* LoF AD risk variants [1], would disrupt lipid metabolism in the mouse brain and alter microglial phenotype on a background of impaired peripheral immune tolerance. To this end, we analyzed brains from three-month-old *Plcg2* wild-type (WT), heterozygous (Het KO), and homozygous knockout (Homo KO) littermates using a multi-omics approach that included shotgun lipidomics, targeted metabolomics, proteomics, and targeted transcriptomics, together with immunofluorescence, and performed flow cytometric profiling of peripheral and brain-draining immune compartments.

## Materials and Methods

In this manuscript, *Plcg2* and *PLCG2* refer to the mouse and human genes, respectively, whereas PLCγ2 refers to the encoded protein.

### Mice, housing, and survival

Experiments were performed on 3-month-old Plcg2 WT, Het KO, and Homo KO mice of both sexes on a congenic C57BL/6J background (MODEL-AD Plcg2 knockout, JAX Strain #029910). All procedures were approved by the Institutional Animal Care and Use Committee at the University of Texas Health Science Center San Antonio and conducted in an AAALAC-accredited, PHS-assured, USDA-registered facility. Mice were housed under specific pathogen-free, Helicobacter-free conditions in an immunocompromised barrier room using sterile individually ventilated sterile cages, with up to five animals per cage, on a 14-h light/10-h dark cycle (lights on at 7:00, off at 21:00) at 24 ± 1.5 °C and a minimum of 10 air changes per hour. Animals had ad libitum access to irradiated pelleted chow (Teklad LM-485 7012) and water.

Our Plcg2 colony was established from three founder breeding cages, each containing one Plcg2 Het KO male and one Plcg2 Het KO female recovered from cryopreserved embryos obtained directly from The Jackson Laboratory. At weaning, tail or ear biopsies were collected for PCR-based genotyping using the Jackson Laboratory protocol, and mice were ear-notched for permanent identification.

For survival analyses, we prospectively recorded litter size at birth and at weaning across more than five years of Plcg2 Het KO × Het KO intercrosses. In total, these crosses generated >650 offspring from 130 litters, with an average of 5.4 pups at birth and 4.7 mice at weaning. Pups that died prior to weaning were counted and, when possible, examined to estimate the timing of death; virtually all observed postnatal deaths occurred between P0 and P5. Genotype distributions among surviving weanlings were used to infer expected versus observed numbers of Plcg2 Homo KO mice under Mendelian inheritance.

### Experimental cohorts and endpoint measurements

The primary cohort comprised 3-month-old Plcg2 WT, Het KO, and Homo KO littermates that did not receive any intervention prior to tissue collection and were used for the endpoint analyses. A second independent cohort consisted of age-matched Plcg2 WT, Het KO, and Homo KO mice that received PBS injections as controls for other experiments; PBS treatment had no detectable effect on body weight, blood glucose, spleen weight, or hemicerebrum weight, so these animals were pooled with the baseline cohort for the expanded analyses (Figure S1 and S3).

At the 3-month endpoint, body weight was measured to the nearest 0.1 g using a calibrated digital scale immediately before euthanasia. Non-fasted blood glucose was measured from tail-vein blood using a Contour glucometer and corresponding test strips (Bayer) between 9:00 and 12:00 with mice maintained on *ad libitum* chow. Mice were then euthanized by isoflurane overdose followed by transcardial perfusion with phosphate-buffered saline (PBS) in accordance with American Veterinary Medical Association guidelines.

After perfusion, brains were hemisected; one hemibrain was post-fixed for histology, and the other was dissected into cerebrum (forebrain minus olfactory bulbs) and brainstem (midbrain, pons, medulla), snap-frozen in liquid nitrogen, and lyophilized at 0.5 mbar for 48 h (Labconco). Lyophilized hemicerebrum samples were weighed on an analytical balance to determine dry hemicerebrum weight. Spleens were dissected, gently blotted to remove excess PBS, and dried at room temperature until constant mass was achieved before weighing on an analytical balance to obtain dry spleen weight.

### Gene Expression Commons (GEXC) hematopoiesis meta-analysis

To contextualize splenic phenotypes, Plcg2 expression across hematopoietic lineages was assessed using the Gene Expression Commons (GEXC) Mouse Hematopoiesis platform, which aggregates 11,939 Affymetrix microarrays spanning hematopoietic stem and progenitor cells, B-cell and T-cell developmental stages, and myeloid lineages. For each probeset, signal intensity was normalized to a common reference to derive a dynamic range of gene expression activity, and Plcg2 expression values were mapped onto a schematic of hematopoietic differentiation to visualize activity across progenitor and mature immune cell compartments (Figure 1E).

**Figure 1.**
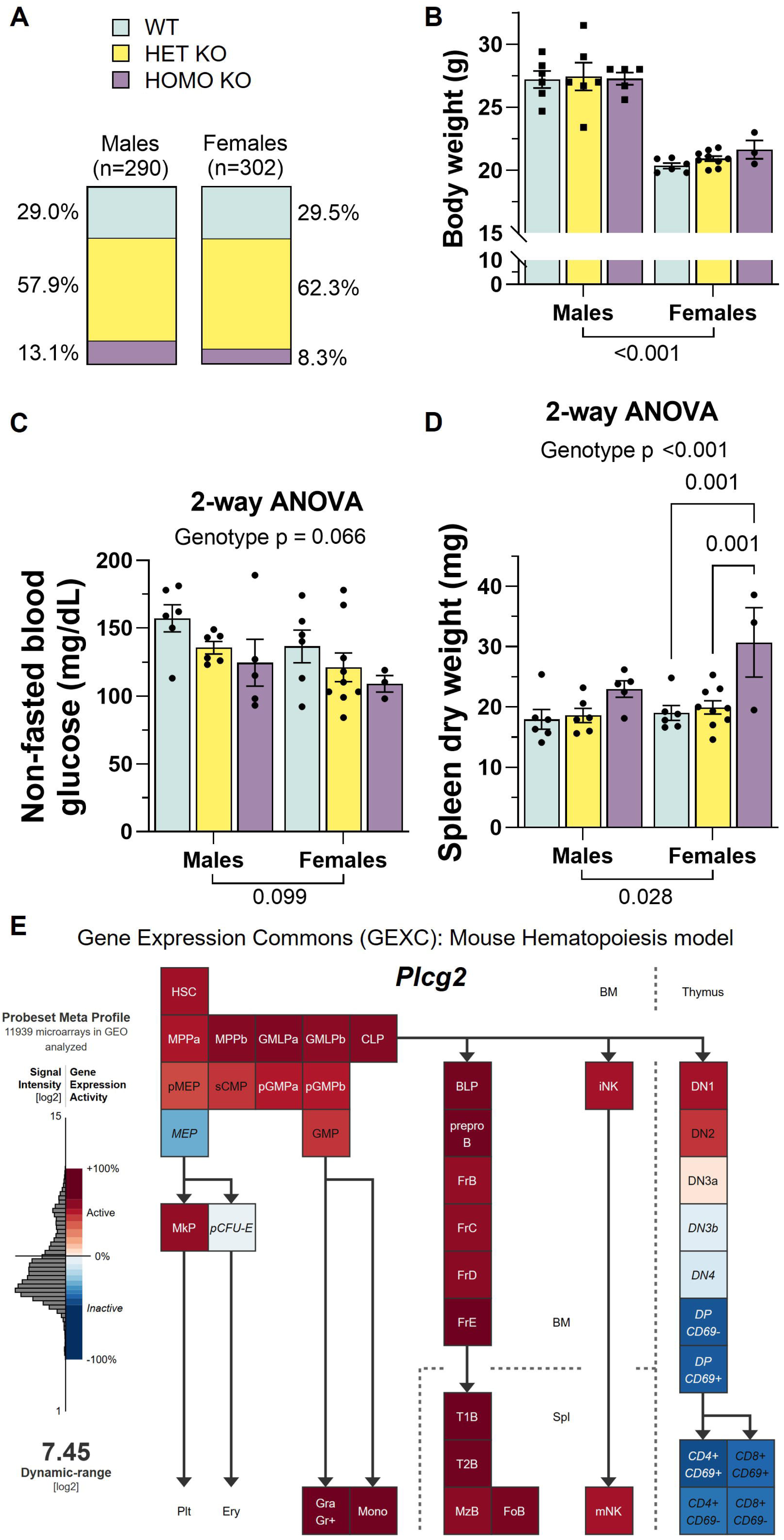
Genotype distribution, body weight, blood glucose, and spleen size in Plcg2-deficient mice. (**A**) Genotype distribution at weaning for male and female offspring from Plcg2 Het KO intercrosses, expressed as the percentage of surviving WT, Het KO, and Homo KO mice. (**B**) Body weight in male and female Plcg2 littermates of the indicated genotypes at the 3-month endpoint. (**C**) Non-fasted blood glucose at the 3-month endpoint in male and female Plcg2 littermates of each genotype. (**D**) Dry spleen weight at the 3-month endpoint in Plcg2 littermates of each genotype. (**E**) Gene Expression Commons (GEXC) Mouse Hematopoiesis meta-profile illustrating *Plcg2* expression across hematopoietic progenitor and mature immune cell populations, including B-cell developmental stages in spleen and blood myeloid progenitor and effector lineages, e.g., granulocyte/macrophage progenitors (GMP), granulocytes (Gra Gr⁺), and monocytes (Mono). Data in B–D are presented as mean ± SEM; individual points represent individual animals (males, squares; females, circles). Two-way ANOVA with factors genotype and sex was followed by Tukey’s post hoc tests assessing simple genotype effects within each sex; p values (adjusted for multiple comparisons) < 0.1 are indicated in the graphs (B-D).

### Shotgun Lipidomics

Cerebrum samples were homogenized in 0.1 PBS using Precellys^®^ ceramic beads/homogenizer equipped with a Cryolys Evolution cooling system (Bertin Technologies). Total protein concentrations were determined using bicinchoninic acid (BCA) protein assay (Thermo Fisher Scientific). Lipids were extracted by a modified procedure of Bligh and Dyer extraction in the presence of internal standards, which were added based on protein content as previously described [44]. Lipids were assessed using a triple-quadrupole mass spectrometer (Thermo Scientific TSQ Altis) and a Quadrupole-Orbitrap™ mass spectrometer (Thermo Q Exactive™) equipped with a Nanomate device (Advion Bioscience Ltd., NY, USA) as previously described [45, 46]. All full and tandem mass spectra were automatically acquired using a customized sequence subroutine operated under Xcalibur software. Data processing including ion peak selection, baseline correction, data transfer, peak intensity comparison, ^13^C deisotoping, and quantitation were conducted using a custom programmed Microsoft Excel macro after considering the principles of lipidomics [47]. The investigator (H.W.) who performed lipid extraction, ran mass spectrometry (MS), and quantified all lipid species was completely blinded to the genotypes of the samples, which were randomized. All samples processed for lipid extraction and MS were included in all analyses (no samples were excluded). Two separate MS runs were performed, one for each sex. Given the high mortality rates of Plcg2 Homo KO females, female samples were run 8 months after the male samples following the exact same methodology and instrumentation. A comprehensive lipidomics data analysis using MetaboAnalyst metadata module with genotype and sex as categorical variables was subsequently performed by unblinded investigators (E.G.K, J.G.R. and J.P.P.). Datasets for classes (18), subclasses (23), and species (188) were log_10_-transformed and analyzed independently. Differentially abundant lipids (DALs) were identified by applying a linear model (limma) based with covariate adjustment (sex), while comparing all genotypes (all contrasts ANOVA-style) and multiple group comparisons using WT or Homo Het as the reference group. FDR-adjusted p value cutoff was 0.05, with 0.1 considered a trend.

### Targeted Metabolomics

Targeted acylcarnitine profiling was performed by the UTMB Mass Spectrometry Facility using its established targeted lipidomics/acylcarnitine panel on a Waters ACQUITY UPLC coupled to a SCIEX QTRAP 6500/7500 triple quadrupole mass spectrometer. Samples were submitted as brain tissue and processed according to the facility’s small-molecule workflow, with extraction performed by MSF staff and quantitative data normalized to tissue weight, as specified on the UTMB small-molecules service request form. The assay employed isotope-labeled surrogate internal standards for each acylcarnitine class, and analytes were quantified from MRM transitions as normalized peak areas relative to the corresponding internal standard signal. The panel included free carnitine (C0), acetylcarnitine (C2), propionylcarnitine (C3), butyrylcarnitine (C4), glutarylcarnitine (C5-DC), hexanoylcarnitine (C6), octanoylcarnitine (C8), decanoylcarnitine (C10), dodecanoylcarnitine (C12), myristoylcarnitine (C14), palmitoylcarnitine (C16), oleoylcarnitine (C18:1), and multiple hydroxylated and unsaturated long-chain acylcarnitines, as reported in the UTMB acylcarnitine panel output file.

For downstream analyses in this study, normalized acylcarnitine areas were exported from the UTMB data matrix, log- or cube-root transformed as appropriate, and analyzed in MetaboAnalyst with sex and Plcg2 genotype specified as categorical variables, using limma-based linear modeling and FDR correction to identify differentially abundant acylcarnitines.

### Nanostring Gene Expression Analysis

NanoString nCounter gene expression analyses quantifies individual RNA molecules without relying on amplification steps. This capability makes NanoString particularly effective at quantifying expression of low and very low abundant genes. Here, we used the Nanostring Glial Profiling Panel to analyze gene expression in cerebrum and brainstem samples. RNA from lyophilized cerebrum and brainstem samples was extracted using the Maxwell^®^ RSC simplyRNA Tissue Kit/Instrument (Promega) with initial homogenization on a TissueLyser LT (Qiagen). RNA concentrations were determined using the Qubit^TM^ RNA Broad Range Assay Kit (Thermo Fisher Scientific). RNA Integrity (RIN), which ranges from 10 (highly intact) to 1 (strongly degraded), was assessed using a TapeStation 4150 via RNA ScreenTapes. 25 ng of RNA per sample (initial concentrations were 50-150 ng/μl with RIN 7-9), loaded into individual NanoString Hybridization reactions for 16 hours at 65°C, and loaded into a NanoString nCounter^®^ SPRINT cartridge/Analysis System. The investigator (S.B.) who performed RNA extractions, hybridization, and ran the SPRING cartridge was blinded to the genotypes of the samples. All RNA samples hybridized and loaded into the SPRINT cartridges were included in all analyses (no samples were excluded). nSolver RCC files were imported into R using the read_rcc function from the *nanostringr* R package by unblinded investigators (H.M. and E.G.K.) **[48]**. Differential gene expression analysis was performed using the *DESeq2* R package **[49]**. Raw NanoString count data were modeled using a negative binomial generalized linear model with the design formula ∼RepeatedAnimal + Sex + Brain.Region + Genotype. RepeatedAnimal accounts for repeated measures from the same animal but different brain region (Brainstem and Cerebrum). Sex and Brain Region were included as covariates. Differential expression for different comparisons such as KO vs WT was assessed using a Wald Test, and p-values were adjusted using the Benjamini-Hochberg procedure. Genes with FDR <0.05 were considered statistically significant. Data was visualized using principal component analysis (PCA) for exploratory data analysis. Volcano plots were generated from DESeq2 differential expression results using custom Python scripts (matplotlib). Statistical significance was determined using Benjamini–Hochberg false discovery rate (FDR) adjusted p-values, with genes meeting an adjusted p-value threshold of <0.05 considered significant. Selected genes were manually annotated according to biological pathways and functional relevance.

### Weighted Gene Co-expression Network Analysis (WGCNA)

RCC files generated from mouse brain samples processed with the NanoString Neuroinflammation Glial Profiling Panel (v1.0; NanoString custom CodeSet), previously used for differential expression analyses, were imported into nSolver Analysis Software v4.0 for downstream weighted gene co-expression network analysis (WGCNA). Sample metadata including sex, genotype, brain region, and cartridge were annotated. All default background correction and normalization parameters were applied, followed by removal of housekeeping genes with CV > 12.5% and average counts < 100. No background thresholding was applied. Only samples passing NanoString QC flags (Imaging, Binding Density, Positive Controls, etc.) were retained. These normalized datasets (mRNA counts) were exported together and separately for male and female animals. To mitigate sex-driven module formation, we separated by sex and conducted WGCNA independently in each group. Sample traits were defined with Genotype as a variable (WT, HET, KO) using only cerebrum samples. Genes with excessive missing values or zero variance were filtered prior to network construction. No additional variance stabilization was applied. A soft-thresholding (b=7) was selected to ensure the resulting network approximated a scale-free topology (scale-free fit index R² > 0.85). Adjacency and topological overlap matrices (TOMs) were generated for network construction. Modules of co expressed genes were identified using hierarchical clustering with the dynamic tree cut algorithm. Modules with similar expression profiles were merged using a cut height of 0.25, corresponding to eigengene correlations ≥ 0.75. Module eigengenes (MEs) were correlated with sample trait (Genotype). Module–trait relationships were evaluated using correlation analysis with FDR correction. Genotype-associated modules were selected for downstream analysis. Intramodular hub “driver” genes were identified using gene significance (GS) and module membership (MM) scores. Gene lists from relevant modules were exported for pathway enrichment using Functional Enrichment analysis within the STRING database.

### Quantitative PCR (qPCR)

RNA purity and quantity was quantified using Cytation 5 with Take3 plate and stored in −80 °C. cDNA was prepared using iScript cDNA synthesis kit (cat 1708891, Bio-Rad) following the manufacturer’s protocol. Gene expression was quantified in triplicate using SYBR green (cat #K0364, Thermo Scientific) for qPCR using a Bio-Rad CFX96 Real-Time PCR detection system. Primers were obtained from Sigma (Supplemental Table 1) and validated to have an efficiency between 80% and 120%. Data were analyzed using the ΔΔCt method with data is represented as relative gene expression compared to WT control mice and actin used as the reference gene.

### Proteomics

Approximately 3 mg of each lyophilized mouse cerebrum sample were placed in a MicroTube fitted with a MicroPestle (Pressure BioSciences), and 30 µl of homogenization buffer [10% SDS in 50 mM triethylammonium bicarbonate (TEAB, Thermo Fisher Scientific) containing protease/phosphatase inhibitors (Halt, Thermo Fisher Scientific) and nuclease (Universal Nuclease, Pierce/Thermo Fisher Scientific). Tissue samples were randomized and homogenized in a Barocycler (Pressure BioSciences, Inc.) for 60 cycles at 35 °C and then the MicroTubes were centrifuged at 21,000 x g for 10 minutes. Aliquots corresponding to 100 µg protein (EZQ™ Protein Quantitation Kit; Thermo Fisher Scientific) were reduced with tris(2-carboxyethyl)phosphine hydrochloride (TCEP), alkylated in the dark with iodoacetamide and applied to S-Traps (mini; Protifi) for tryptic digestion (sequencing grade; Promega) in 50 mM TEAB. Peptides were eluted from the S-Traps with 0.2% formic acid in 50% aqueous acetonitrile and quantified using Pierce™ Quantitative Fluorometric Peptide Assay (Thermo Fisher Scientific).

Data-independent acquisition mass spectrometry was conducted on an Orbitrap Fusion Lumos mass spectrometer (Thermo Fisher Scientific). On-line HPLC separation was accomplished with an RSLC NANO HPLC system (Thermo Fisher Scientific/Dyonex). A pool was made of all of the samples, and 2-µg peptide aliquots were analyzed using gas-phase fractionation with staggered 4-m/z windows (30k resolution for precursor and product ion scans, all in the orbitrap). The 4-mz data files were used to create a DIA chromatogram library **[50]** by searching against a Prosit-generated predicted spectral library **[51]** based on the UniProt_Mus musculus reviewed library_20191022. Mass spectrometry data for experimental samples were acquired in the Orbitrap using 8-m/z windows and searched against the chromatogram library. Mass spectrometry data was processed in Scaffold DIA (Proteome Software) filtered at 1% FDR using a minimum peptide count of 2. The investigators (S.P. and S.W.) who processed the samples and ran MS data were blinded to the genotypes of the samples. Two separate MS runs were performed, one for each sex. Given the high mortality rates of Plcg2 Homo KO females and funding limitations, female samples were run more almost two years after male samples were run following the exact same methodology, instrumentation, and reference database.

A comprehensive proteomics data analysis using MetaboAnalyst metadata module with genotype and sex as categorical variables was subsequently performed by unblinded investigators (G.C., E.G.K. and J.P.P.). Log_10_ normalized non-scaled abundances of the 5,501 identified proteins that passed the filters mentioned above (Fig. S2A) were imported. For preprocessing, coefficients of variation (CVs) were calculated across all proteins. To reduce the influence of highly variable measurements, proteins exceeding the 95th percentile of CV values were excluded. Among the initial 5,501 detected proteins, the 95th percentile CV threshold was 32.449, resulting in the removal of the top 5% most variable proteins leaving 5,225 proteins for downstream analysis. Because samples were acquired across two independent mass spectrometry runs, an additional filtering step was performed to minimize potential batch-driven artifacts related to unequal protein detection between sexes. Proteins that were completely absent in all samples from either males or females were removed, as these missing values were considered more likely to reflect technical differences between runs rather than true biological effects. Specifically, 422 proteins lacked detectable values in all male samples, while 325 proteins lacked detectable values in all female samples. After applying both the CV-based filtering and unequal-presence filtering steps, a final set of 4,478 proteins was retained for downstream analyses and wrangled using MetaboAnalyst Data display outcomes included metadata overview (Fig. 8A), PCA (Fig. S2B), interactive heatmap (Fig. 8C). Differentially abundant proteins (DAPs) were identified by applying a linear model based on limma using WT as the reference group, all contrasts ANOVA-style, and unspecified blocking factor. FDR-adjusted p value cutoff was 0.05. KEGG enrichment analysis was performed using Enrichr (Fig. 8E) and a functional protein association network was performed using STRING v12.0 (Fig. 8E) using the 25 identified DAPs. Finally, a secondary PCA was performed using only the 575 proteins that had a raw p value below 0.05 (Fig. 6E).

### Immunofluorescent staining

Sections were permeabilized in 0.5% Triton, blocked in 10% normal goat serum, and incubated with primary antibodies overnight at 4°C (pan-myeloid marker ionized calcium binding adaptor protein 1 (Iba1): WAKO, neuronal nuclei (NeuN): A60, EMD Millipore, myelin associated glycoprotein (MAG): D4G3, Cell Signaling, glial fibrillary acidic protein (GFAP): GA5, eBiosceince Invitrogen), P2RY12: Synaptic Systems 476 308), myelin basic protein (MBP): Invitrogen PA1-10008 and with secondary antibodies for 90 minutes (Invitrogen goat anti-rabbit AlexaFluor 555, goat anti-mouse AlexaFluor 488, goat anti chicken, goat anti guinea pig). Sections were mounted on permafrost slides, counterstained with DAPI, coverslips mounted and sealed. Slides were imaged using a Bio-Tek Cytation 5 multi-mode imager. For image analysis, one region of interest (ROI) inclusive of forebrain minus olfactory lobe (as in Figure 7C) per animal was selected and an ImageJ image processing analysis macro used to threshold the image and generate the area coverage.

### Flow cytometry

Upon euthanasia, cervical lymph nodes and spleens were removed and placed immediately in ice-cold medium (Hank’s balanced salt solution, 5% FBS, 10 mg/ml DNAse.) All subsequent steps were performed at 4°C. The tissues were lacerated with a scalpel and pressed through a wire mesh using the plunger from a 3 ml syringe. Red blood cells were lysed in spleen samples using ACK lysis buffer (Thermo Fisher Scientific, Waltham, MA, USA). Cell suspensions were counted using a hemacytometer. Samples were stained with LIVE/DEAD Blue (Invitrogen, catalog no. L23105; dilution 1:1000). For CD4 Treg analyses, samples were incubated with a cocktail of fluorescently labeled antibodies containing APC-Cy-7-conjugated anti-CD4 (Clone RM4-5; Biolegend; dilution 1:150), BV 395-conjugated anti-CD8 (Clone 53-6.7; BD Horizon; dilution 1:200), PE-conjugated anti-CD25 (Clone PC61; BD Biosciences; dilution 1:200), and BV650-conjugated anti-CD44 (Clone IM7; Biolegend; dilution 1:200). Following treatment with Foxp3 fixation/permeabilization buffer (eBioscience, catalog no. 00-5523-00), samples were stained with PE-Cy7-conjugated anti-Foxp3 (Clone FJK-16s; eBioscience; dilution 1:250). For B cell analysis, single cell suspensions were stained with BV510-conjugated anti-CD19 (Clone: 1D3; BD Biosciences; dilution 1:100), and after fixation and permeabilization as described for CD4 Tregs, BV711-conjugated anti-T-bet (Clone: 4B10; Biolegend; dilution 1:100). For CD8 Treg analysis, single cell suspensions were stained with PE-Cy7-conjugated anti-CD122 (Clone: TM-b1; Biolegend; dilution 1:100), APC-Cy-7-conjugated anti-CD8a (Clone: 53-6.7; BD Pharmingen; dilution 1:100), APC-conjugated anti-CD3e (Clone: 145-2C11; BD Bioscience; dilution 1:100), and PE-conjugated anti-Ly49, (Clone: 14B11; Biolegend; dilution 1:200). For brain analysis, single cell suspensions were generated using enzyme dissociation (Adult Brain Dissociation Kit, Miltenyi Biotec) as previously reported [52]. Samples with stained with an antibody cocktail containing APC-conjugated anti-CD45.2 (Clone: 104; eBioscience; dilution 1:100), BV510-conjugated anti-CD11c (Clone: N4181; Biolegend; dilution 1:100), and Alexa Fluor488-conjugated anti-CD11b (Clone: M1/70, Biolegend, dilution 1:100). DAPI staining was used to exclude dead/dying cells in brain samples. Flow cytometry analysis was performed on the Cytek Aurora spectral flow cytometer running SpectroFlo v3.3.0. Data were analyzed using FlowJo software (v10.9, Tree Star).

### Statistics

In general, univariate data were analyzed in GraphPad Prism using one-way (genotype) or two-way (genotype × sex) analysis of variance (ANOVA), followed by Tukey’s post hoc tests to control for multiple comparisons. Specific statistical tests and p values for each graph are provided in the corresponding figure legends. Multivariate datasets were analyzed using method-specific approaches as outlined above (e.g., limma for lipidomics and proteomics, DESeq2 for NanoString, WGCNA for co-expression networks). A p value between 0.05 and 0.10 was considered a trend, and p < 0.05 was considered statistically significant.

## Results

### *Pclg2* deficiency substantially impairs early survival and modestly alters basal glucose homeostasis

Across more than five years of colony maintenance under specific pathogen-free, *Helicobacter*-free barrier conditions under sterile housing conditions commonly used for immunocompromised mouse strains, Plcg2 Het intercrosses generated >650 offspring from 130 litters, with an average of 5.4 pups at birth and 4.7 mice at weaning, indicating that approximately one pup every two litters died postnatally prior to weaning (typically between P0-P5 by direct observation). Under Mendelian inheritance, 25% of weaned mice of each sex are expected to be Plcg2 Homo KO, yet only 13.1% of surviving males and 8.3% of surviving females were Homo KO at weaning, implying that roughly half of Homo KO males and three-quarters of Homo KO females died before weaning (Figure 1A). Across litters, 60 pups died postnatally, accounting for only about half of the ∼115 deaths predicted from Mendelian ratios, indicating that approximately half of Plcg2 Homo KO mice die *in utero*.

Despite elevated pre-weaning mortality, all Plcg2 Homo KO mice that survived to weaning lived to the planned three-month endpoint and displayed normal body weight relative to WT and Het KO littermates. Females weighed less than males at 3 months (23% lower mean body weight), but within each sex there were no significant differences among genotypes (Figure 1B and Figure S1A). To assess basal metabolic status, we evaluated non-fasted glucose at 3 months and observed a trend toward a genotype main effect at 3 months (Figure 1C). When saline-treated controls from an independent cohort were included, this effect reached statistical significance, with lower glucose in Homo KO mice and Het KO animals exhibiting intermediate values, non-fasted glucose was reduced by 15% in males and 11% in females relative to WT, with statistically significant decreases in males (Figure S1B).

### *Plcg2* deficiency induces myeloid-skewed splenomegaly and coordinated ABC/Treg imbalance

Dry spleen weight was significantly increased in Plcg2 Homo KO mice compared with WT and Het KO littermates at the 3-month endpoint (Figure 1D). Inclusion of saline-treated controls confirmed robust splenomegaly in Homo KO mice, primarily driven by females (48% increase) with similar directional trends in males (19% increase; Figure S1C).

To determine whether splenomegaly reflected global or subset-specific changes in spleen immune cell composition, we first quantified total live splenocytes. Total spleen cellularity was not significantly different between Plcg2 Homo KO and WT mice (Figure S2A), indicating that enlarged spleens are not explained by a general increase in total spleen cell number. We then examined two broad spleen immune cell compartments: CD19⁻CD11b⁺ cells, a population enriched for myeloid/innate cells (e.g. monocytes/macrophages and granulocytes), and CD19⁺CD11b⁻ B cells (Figure 2A–B), quantified both as absolute cell numbers and as percentages of total spleen cells. Plcg2 Homo KO spleens exhibited a 2.2-fold increase in the proportion of CD19⁻CD11b⁺ cells and a 39% decrease in CD19⁺CD11b⁻ B cells (Figure 2C–D) relative to WT littermates, indicating a myeloid-skewed spleen cell composition in the absence of PLCγ2. Similar directional changes were observed when analyzing absolute cell numbers (Figure S2B-C).

**Figure 2.**
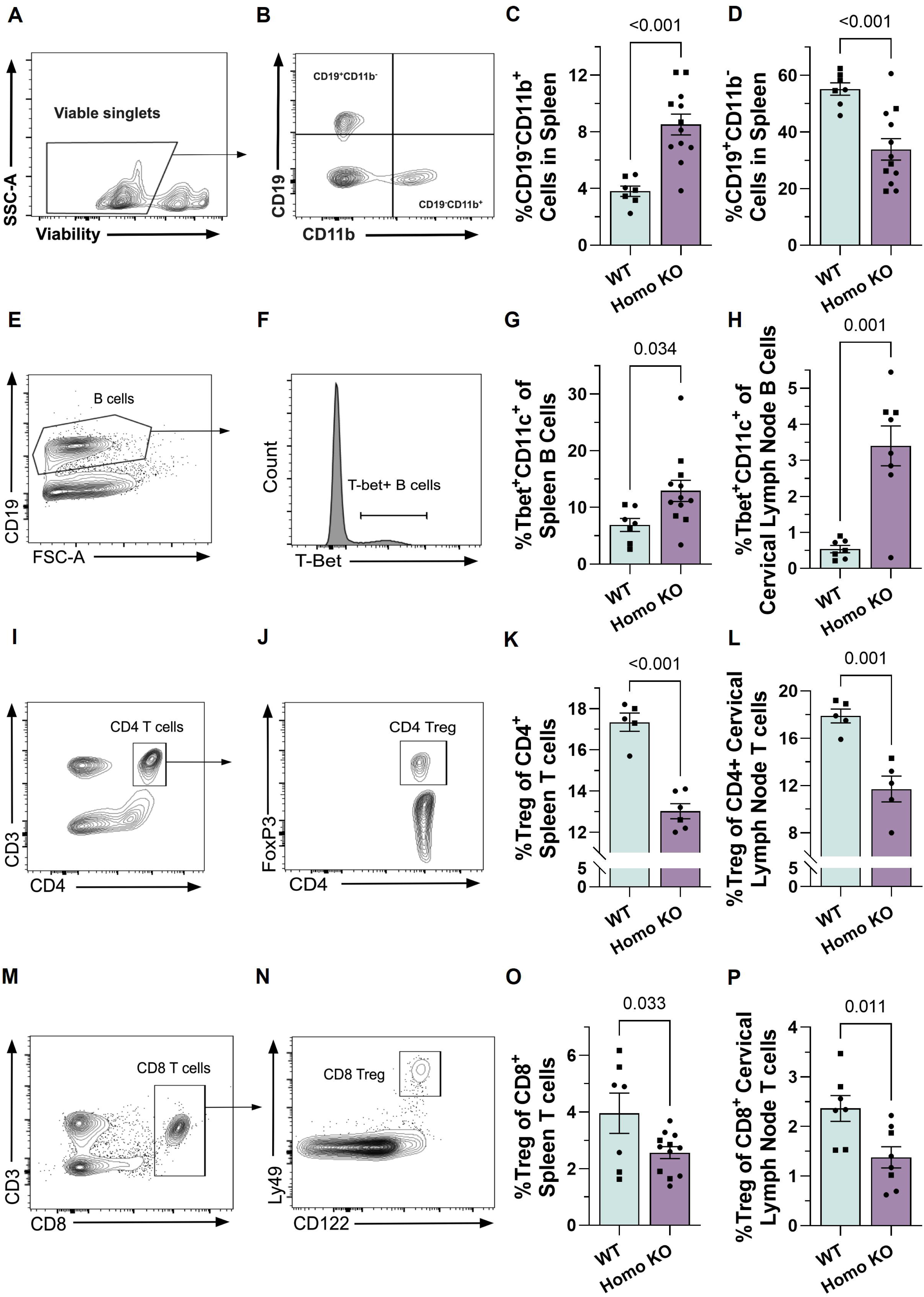
Myeloid/B-cell composition, age-associated B cells, and regulatory T cells in spleen and cervical lymph nodes from Plcg2-deficient mice. (**A-B**) Representative flow cytometry gating strategy for CD19⁻CD11b⁺ spleen cells. (**C**) Percentage of CD19⁻CD11b⁺ among total spleen cells. (**D**) Percentage of CD19⁺CD11b⁻ B cells among total spleen cells. (**E**-**F**) Representative gating strategies for CD19⁺ B cells and ABC-like (CD11c⁺T-bet⁺) B cells within CD19⁺ B cells. (**G**) Percentage of ABC-like (CD11c⁺T-bet⁺) CD19⁺ B cells in spleen. (**H**) Percentage of ABC-like (CD11c⁺T-bet⁺) CD19⁺ B cells in cervical lymph nodes. (**I**-**J**) Representative gating strategies for CD4⁺ T cells and FoxP3⁺ CD4 Tregs. (**K**) Percentage of FoxP3⁺ CD4 Tregs among CD4⁺ T cells in spleen. (**L**) Percentage of FoxP3⁺ CD4 Tregs among CD4⁺ T cells in cervical lymph nodes. (**M**-**N**) Representative gating strategies for CD8⁺ T cells and CD8 Tregs (Ly49⁺CD122⁺). (**O**) Percentage of CD8 Tregs among CD8⁺ T cells in spleen. (**P**) Percentage of CD8 Tregs among CD8⁺ T cells in cervical lymph nodes. ABC-like (CD11c⁺T-bet⁺) CD19⁺ B cells were quantified in spleen and cervical lymph nodes. Data are shown as representative dot plots (mean ± SEM); individual points represent individual animals (males, squares; females, circles). Samples with insufficient cell yield for reliable gating were excluded a priori. Statistical significance was assessed using unpaired two-tailed t tests when data passed normality and homoscedasticity tests (D, G, K–L, P) and Welch’s t tests when F tests indicated unequal variances between groups (B, H, O); p values < 0.1 are indicated in the graphs.

Within the remaining CD19⁺ B-cell compartment, and given that PLCγ2 functions immediately downstream of the B-cell receptor (BCR) [53], is critical for BCR-induced calcium signaling, and is linked with longevity, we hypothesized that Plcg2 loss might perturb age-associated B-cell (ABC) populations. ABCs are defined by expression of CD11c and the transcription factor T-bet and arise in response to chronic BCR and innate stimulation [54]. We therefore quantified ABC-like (CD11c⁺T-bet⁺) B cells in spleen as a proportion of total CD19^+^ B cells using a sequential gating strategy (FSC/SSC, singlets, viable cells, CD19^+^ B cells, CD11c^+^T-bet^+^ cells; Figure 2C-D). ABC-like B cells were significantly increased in Plcg2 Homo KO spleen (87% increase; Figure 2E), indicating enhanced accumulation of this age-associated subset in a major peripheral lymphoid organ.

Because cervical lymph nodes drain the head and neck and are positioned to integrate immune signals arising from the meninges and glymphatic outflow [55], we next examined ABC-like B cells in this brain-draining compartment. Plcg2 Homo KO mice exhibited markedly increased frequencies of T-bet⁺ (ABC-like) B cells in cervical lymph nodes (6.3-fold increase; Figure 2F), demonstrating that age-associated B-cell expansion extends to lymphoid tissues that interface directly with the central nervous system.

Given these B-cell changes and the central role of regulatory T cells in restraining age- and autoimmunity-associated inflammation [56], we next assessed CD4^+^ and CD8^+^ T-cell populations. FoxP3⁺ CD4 regulatory T cells (Tregs), a key regulatory subset that maintains peripheral tolerance, were significantly reduced among CD4⁺ T cells in Plcg2 Homo KO spleen (25% decrease; Figure 2E). Similar reductions in FoxP3⁺ CD4 Tregs were detected in cervical lymph nodes (35% decrease; Figure 2F). CD8 Tregs, a non-redundant regulatory subset, were quantified within live CD8⁺ T cells using FSC/SSC, singlet, viability, CD3⁺CD8⁺ gating followed by Ly49 and CD122 expression (Figure 2G–H). Plcg2 Homo KO mice exhibited significantly lower frequencies of CD8 Tregs in spleen (35% decrease) and cervical lymph nodes (42% decrease, Figure 2I–J), indicating that CD4 and CD8 regulatory T-cell compartments are diminished in peripheral and brain-draining lymphoid tissues in the absence of PLCγ2 expression.

### *Plcg2* expression across neuroimmune compartments

To determine whether *Plcg2* deficiency impacts brain structure at three months of age, we measured dry cerebrum weight and observed no differences across genotypes, indicating no gross brain atrophy in this young adult time window in either sex (Figure 3A and Figure S1D). Flow cytometric analysis of dissociated brain tissue revealed that CD19⁺ B cells and CD3⁺ T cells were very rare and thus near the limit of reliable detection, precluding robust quantification of these populations. In contrast, microglia were readily detectable, allowing us to assess expression of the activation marker CD11c (*Itgax*) within the microglial gate (FSC/SSC, singlet, viability, CD11b⁺CD45^mid^ gating; Figure 3B). Plcg2 Homo KO mice showed a significant increase of 10% in CD11c mean fluorescence intensity (MFI) in microglia compared to WT littermates (Figure 3C-D), indicating a modest shift toward a more CD11c-high microglial phenotype at this age.

**Figure 3.**
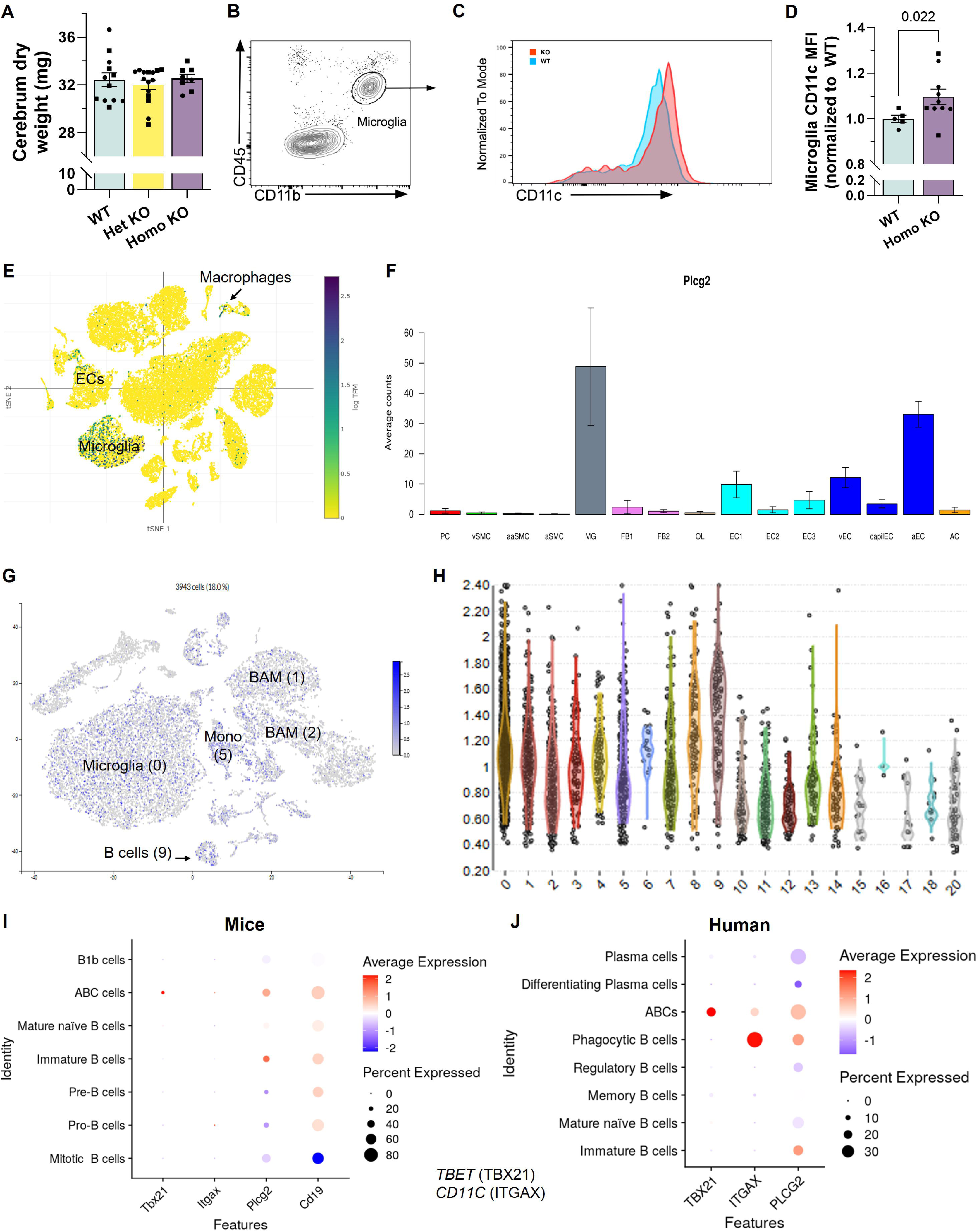
Microglial activation-associated marker abundance in Plcg2-deficient mice and Plcg2 transcript distribution across neuroimmune and vascular compartments. (**A**) Cerebrum dry weight in three-month-old Plcg2 WT, Het KO, and Homo KO mice to assess gross brain structure. (**B**) Flow cytometry gating strategy used to identify microglia in dissociated brain tissue. (**C**-**D**) Abundance of the activation-associated surface marker CD11c, quantified as mean fluorescence intensity (MFI) within the microglial gate in Plcg2 WT and Homo KO mice and normalized to WT. (**E**-**H**) Publicly available mouse single-cell RNA-seq datasets from WT brains were analyzed to visualize Plcg2 expression across brain cell types, including microglia, macrophages, endothelial cells, and other neural and vascular populations. (**I**-**J**) Additional mouse and human single-cell B-cell atlases were used to examine Plcg2/PLCG2 expression in ABC-like and other B-cell subsets at CNS borders. Data are presented as bar graphs, density plots, dimensionality-reduction maps, violin plots, and dot plots as indicated. Statistical analysis was performed using one-way ANOVA with Tukey’s post hoc tests for cerebrum dry weight (A) and unpaired two-tailed t tests for microglial CD11c MFI (D); data are shown as representative dot plots (mean ± SEM); individual points represent individual animals (males, squares; females, circles).

To contextualize these findings, we queried public mouse and human single-cell and single-nucleus RNA-seq atlases to define the cellular distribution of *Plcg2/PLCG2* in brain and brain-border tissues. Across datasets, *Plcg2* transcripts were enriched in microglia and brain-associated macrophages, with additional expression in endothelial and perivascular macrophage-like cells and minimal expression in neurons, astrocytes, and mature oligodendrocytes (Figure 3E-F).

In brain and meningeal immune atlases, *Plcg2* was detected in a substantial fraction of CD45⁺ cells, including parenchymal microglia, multiple border-associated macrophage subsets at choroid plexus and dural surfaces (Figure 3H), and ABC-like B-cell clusters in meningeal and brain-associated lymphoid tissues (Figure 3I). Similar PLCG2-high B-cell populations were present in human post-mortem brain immune datasets (Figure 3J).

### Plcg2 deficiency produces subtle, multi-omic changes in the brain with prominent lipid alterations

Having established that *Plcg2* is selectively expressed in microglia, BAMs, border B cells, and endothelial cells at CNS interfaces, and given the strong genetic association between *PLCG2* and AD, we next asked how loss of Plcg2 affects global brain molecular profiles using an unbiased multi-omics approach. Cerebral tissues (forebrain with olfactory bulbs removed) from Plcg2 WT, Het KO, and Homo KO mice were lyophilized and pulverized, and powdered material was split for mass spectrometry-based proteomics, NanoString nCounter Glial Profiling of RNA, and multidimensional mass spectrometry-based shotgun lipidomics. All omics assays were performed in sex-stratified batches (separate male and female cohorts). All available cerebrum samples were analyzed by proteomics and lipidomics, whereas RNA profiling was performed on a subset of cerebrum samples (due to the 12-lane capacity of each NanoString cartridge) as well as on brainstem samples. Global lipidomic profiling demonstrated the most pronounced effect of *Plcg2* loss. After quality filtering, we quantified close to 200 brain lipid species in total for each sex, after excluding species with high within-group variability (coefficient of variation >40%) and additional species that failed our internal curation rules, which were based on established principles of lipidomics [44]. We then fit a linear model adjusting for sex, restricted to 188 lipid species reliably quantified in both male and female runs. Principal component analysis (PCA) revealed clear genotype-associated separation of Plcg2 homozygous knockout samples from WT and Het KO brains in both sexes, with significant PERMANOVAs results for each sex (Figure 4A–B). Volcano-plot analysis identified a focused set of 17 differentially abundant lipids between Plcg2 Homo KO and WT brains; strikingly, 16 of these species were decreased and only 1 was increased, yielding a predominantly loss-of-function signature. These under-abundant lipids were enriched in myelin and phosphoinositide pathways, including multiple hexosylceramide (cerebroside) and phosphatidylinositol-4,5-bisphosphate (PIP_2_) species in Plcg2-deficient mice (Figure 4C). Although our shotgun lipidomics methodology did not distinguish galactocerebrosides from glucocerebrosides, the affected cerebrosides included very-long-chain molecular forms that are almost exclusively myelin-associated, and prior work indicates that the overwhelming majority of brain hexosylceramides are galactosylceramides (galactocerebrosides) rather than glucocerebrosides [57, 58]. In parallel, the differentially abundant phosphoinositides comprised PIP_2_ species, the canonical substrate of PLCγ2.

**Figure 4.**
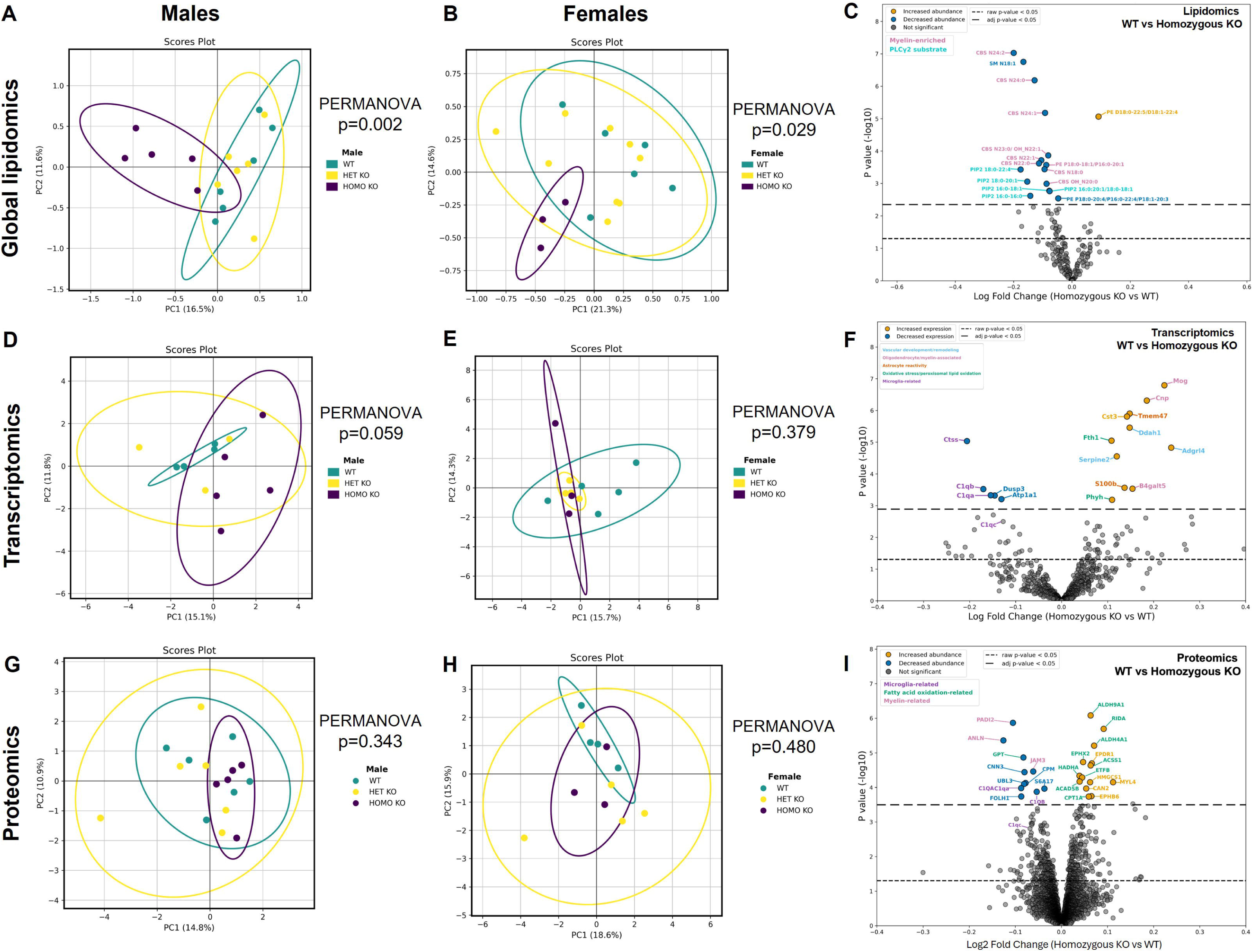
Multi-omic clustering and differential-abundance analysis of Plcg2-deficient brains across lipid, RNA, and protein layers. (**A**-**B**) Principal component analysis (PCA) of global brain lipidomic profiles is shown for male and female Plcg2 WT, Het KO, and Homo KO mice, with 95% confidence ellipses and PERMANOVA p values indicating genotype-associated dispersion in each sex. (**C**) Volcano plots depict differentially abundant brain lipids between Plcg2 Homo KO and WT mice, highlighting myelin-associated and phosphoinositide species. (**D**-**E**) PCA of NanoString glial transcriptomic profiles illustrates genotype clustering patterns in male and female brains, with corresponding PERMANOVA results. (**F**) Volcano plot summarizing differentially expressed genes between Plcg2 Homo KO and WT brains. (**G**-**H**) PCA of proteomic data from male and female brains shows global protein-level dispersion across Plcg2 genotypes. (**I**) Volcano plot displaying differentially abundant proteins between Plcg2 Homo KO and WT brains. Linear models and multiple-testing corrections used for each omic layer are described in the Methods.

Transcriptomic profiling with the NanoString nCounter Glial Profiling Panel, which measures expression of 770 mouse genes across 55 pathways spanning five major themes (cell stress and damage responses, glial homeostasis and activation, inflammation and peripheral immune invasion, neurotransmission, and signaling pathways that regulate glia), revealed more modest genotype effects. PCA showed a modest trend toward separation of *Plcg2* KO samples from WT and Het KO clusters in males, with PERMANOVA results that were borderline non-significant (Figure 4D), whereas females displayed substantial overlap among *Plcg2* genotypes with non-significant PERMANOVA results (Figure 4E). Consistent with this limited separation, differential expression analysis using DESeq2 with sex and brain region included as covariates identified a small set of 16 differentially expressed genes in *Plcg2* Homo KO versus WT brains, with mild fold-change magnitudes, including 5 downregulated and 11 upregulated transcripts (Figure 4F). Most downregulated genes were microglia-enriched, whereas upregulated genes were associated with myelin/oligodendrocyte pathways, consistent with the global lipidomics data, and, unexpectedly, with fatty-acid oxidation and vascular functions. Proteomic profiling showed the smallest global impact of *Plcg2* loss. Because batch effects were substantial and could not be clearly distinguished from true sex differences, downstream analyses were restricted to proteins detected in both runs after excluding features with high intra-group variability (coefficient of variation >32%, 5% of quantified proteins), yielding 4,478 proteins for analysis from an initial set of >5,400 quantified proteins. In both sexes, brain proteomic profiles showed only weak genotype-associated dispersion with largely overlapping 95% confidence ellipses in PCA space with no clear separation of Plcg2 genotypes by PERMANOVA (Figure 4G–H). Nonetheless, differential abundance analysis using a linear model with sex as a covariate identified a small set of 24 proteins altered in *Plcg2* Homo KO versus WT brains, with very modest fold-change magnitudes. Of these, 11 were decreased and 13 increased (Figure 4F); decreased proteins were dominated by microglia-enriched complement components, whereas increased proteins included multiple enzymes involved in fatty-acid oxidation.

Together, these multi-omic data indicate that Plcg2 deficiency exerts its strongest impact on brain lipid metabolism, particularly myelin-associated and PIP_2_ species, with more modest effects on glial RNA signatures and minimal global-scale alterations in the proteome, while still showing discrete protein-level changes that converge on microglial complement pathways and fatty-acid oxidation.

### Co-expression modules link *Plcg2* genotype to immune, metabolic, and myelin gene networks

To determine whether the modest transcriptomic changes detected by NanoString coalesced into coherent gene networks associated with *Plcg2* genotype, we applied weighted gene co-expression network analysis (WGCNA) to cerebrum expression data. The cerebrum was selected because it was available from both sexes and because strong differences between cerebrum and brainstem dominated when regions were analyzed together. Since male and female samples were processed on separate cartridges and initial analyses indicated that mixed-sex networks were driven largely by sex/run effects, WGCNA was performed separately within each sex. This analysis identified several modules whose eigengenes correlated with *Plcg2* genotype in a sex-dependent manner, including two modules positively associated with *Plcg2* Homo KO status (one in males and one in females) and one module negatively associated with genotype in females (Figure 5A).

**Figure 5.**
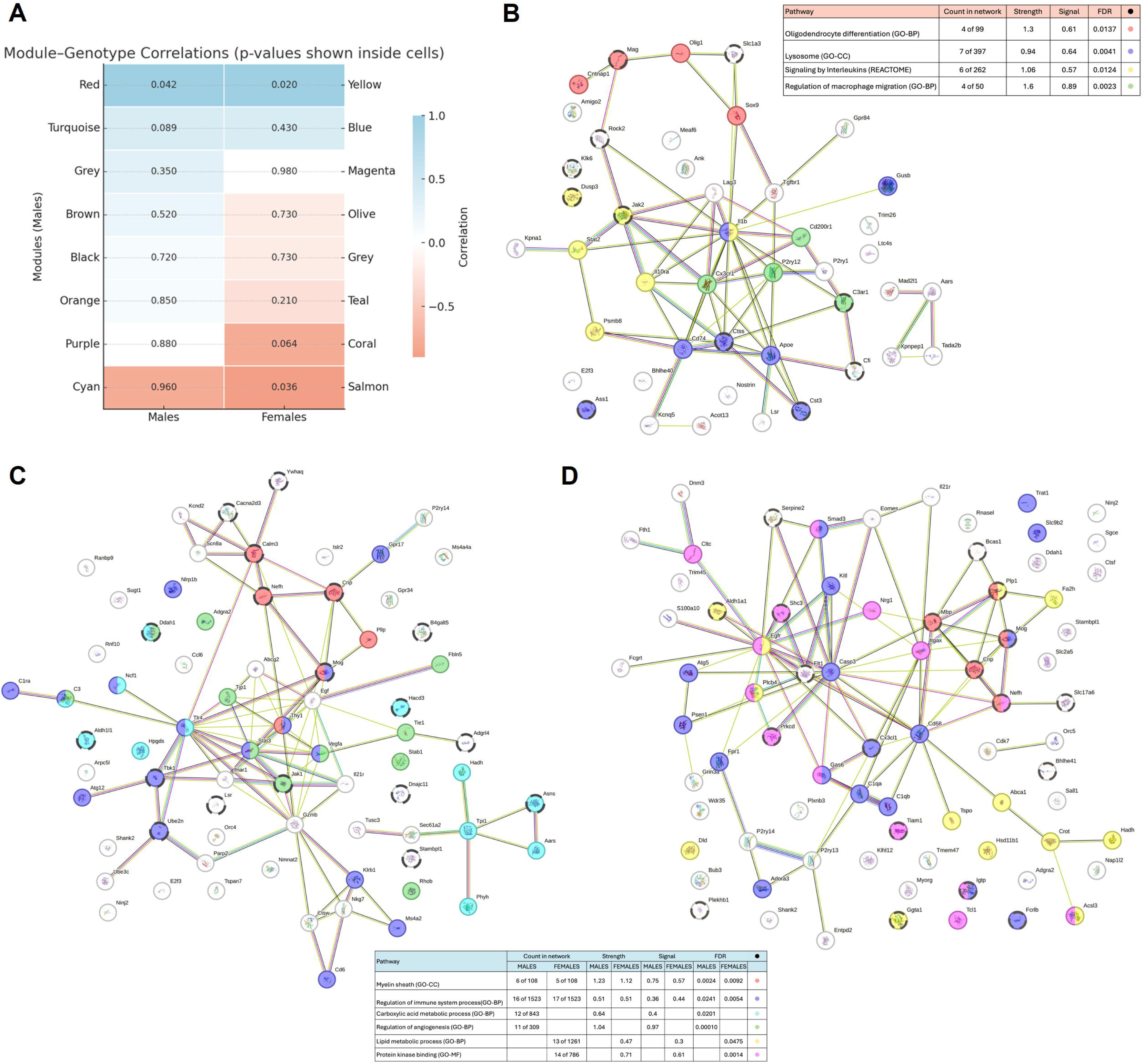
WGCNA modules linking Plcg2 genotype to immune, metabolic, and myelin gene networks. Weighted gene co-expression network analysis (WGCNA) was performed on NanoString glial panel data from cerebrum, with male and female samples analyzed in separate sex-specific networks. (**A**) Module–genotype correlations are shown as heatmaps of module eigengenes versus Plcg2 genotype for males and females, with correlation coefficients and corresponding p values indicated within each cell. (**B**) Gene ontology and pathway enrichment analysis for the female-biased negatively correlated module highlights FDR-significant terms related to oligodendrocyte differentiation, lysosomal function, regulation of macrophage migration, and signaling by interleukins. (**C**-**D**) Enrichment analysis for male- and female-biased positively correlated modules shows FDR-significant over-representation of myelin sheath, immune regulation, lipid and carboxylic-acid metabolic processes, angiogenesis, and protein kinase binding pathways. Data are presented as module–trait correlation heatmaps and pathway-enrichment plots; WGCNA parameters, module-definition criteria, and enrichment analysis methods are detailed in the Methods.

**Figure 6.**
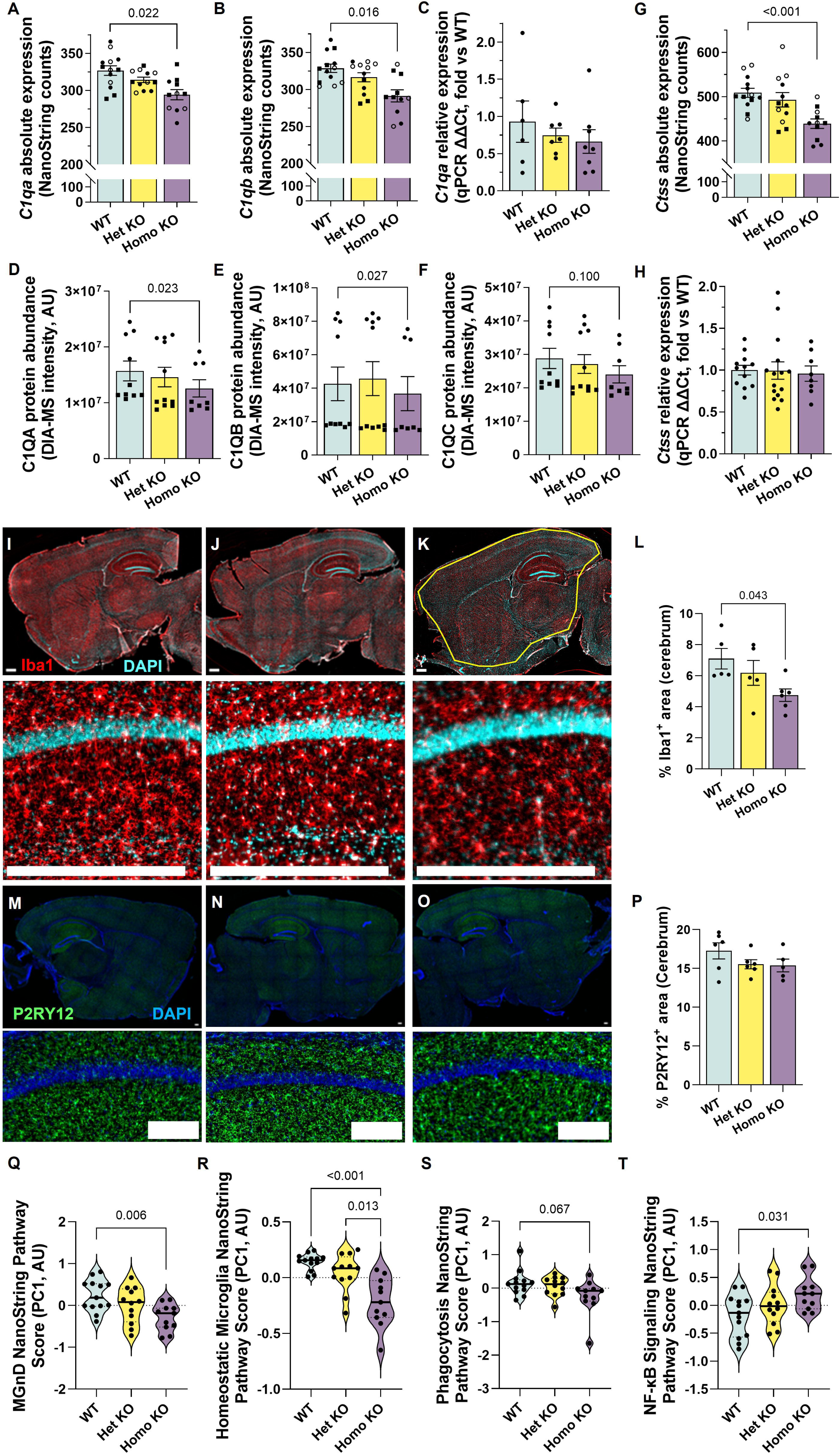
Multi-level assessment of microglial complement, lysosomal markers, morphology, and pathway scores in Plcg2-deficient brains. (**A**-**B**, **G**) C1qa, C1qb, and Ctss transcripts were quantified in cerebrum and brainstem using the NanoString Glial Profiling Panel, with absolute NanoString counts shown for each genotype. (**C**, **H**) C1qa and Ctss mRNA levels were additionally measured in cerebrum by qPCR and expressed relative to WT using the ΔΔCt method. (**D**-**F**) DIA-MS proteomics was used to quantify C1QA, C1QB, and C1QC protein abundance in cerebrum, shown as normalized intensities for Plcg2 WT, Het KO, and Homo KO mice. Immunofluorescence staining of cerebrum sections for Iba1 and P2RY12 was used to assess microglial coverage, with representative images and quantification of Iba1⁺ and P2RY12⁺ area fractions shown for each genotype (**I**-**P**). NanoString-derived pathway scores based on curated microglial gene sets summarize neurodegeneration-associated (MgND), homeostatic, phagocytosis, and NF-κB signaling programs across Plcg2 genotypes (**Q**-**T**). Data are presented as bar plots, violin plots, and representative images, with each point representing an individual animal; covariates and linear models used for NanoString and proteomics data are described in the Methods. Statistical analysis was performed using ordinary one-way ANOVA with Tukey’s post hoc tests for panels L-T, except for panel R where a Welch’s ANOVA with Dunnett’s T3 post hoc test was used after Brown-Forsythe and Bartlett’s tests indicated unequal variances between groups.

**Figure 7.**
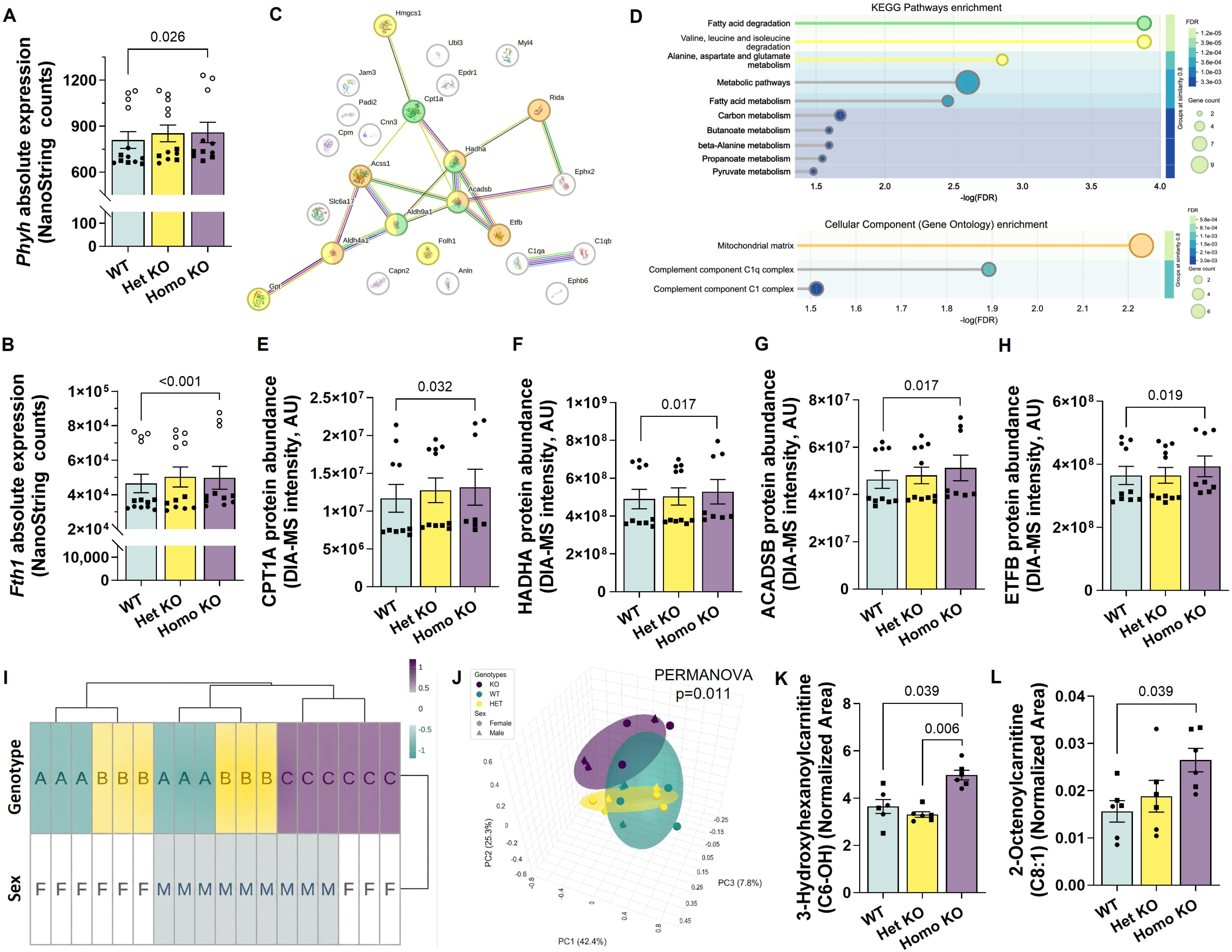
Mitochondrial carboxylic- and fatty-acid oxidation pathways and acylcarnitine profiles in Plcg2-deficient brains. (**A**-**B**) Phyh and Fth1 transcripts were quantified in cerebrum using the NanoString Glial Profiling Panel, with absolute NanoString counts shown for Plcg2 WT, Het KO, and Homo KO mice. (**C**-**D**) A STRING protein–protein interaction network illustrates differentially abundant proteins associated with Plcg2 genotype, and bubble plots summarize KEGG pathway and Gene Ontology cellular-component enrichment, including fatty-acid and amino-acid metabolic pathways and mitochondrial matrix localization. (**E**-**H**) DIA-MS proteomics was used to quantify the fatty-acid β-oxidation enzymes CPT1A and HADHA and the amino-acid/branched-chain acyl-CoA dehydrogenase ACADSB, as well as the electron transfer flavoprotein subunit ETFB, with normalized protein abundances shown for each genotype. (**I**-**J**) Targeted LC/MS acylcarnitine metabolomics was performed on cerebrum extracts from an independent cohort of Plcg2 WT, Het KO, and Homo KO mice, and a metadata heatmap and PCA plot illustrate genotype- and sex-associated clustering patterns. (**K**-**L**) Normalized levels of selected acylcarnitine species, including 3-hydroxyhexanoylcarnitine (C6-OH) and 2-octenoylcarnitine (C8:1), are shown for each genotype. Data are presented as bar plots, STRING interaction maps, enrichment bubble plots, heatmaps, and PCA score plots, with each point in bar plots representing an individual animal; statistical models, covariates, and multiple-testing corrections for NanoString, proteomics, and metabolomics analyses are detailed in the Methods.

**Figure 8.**
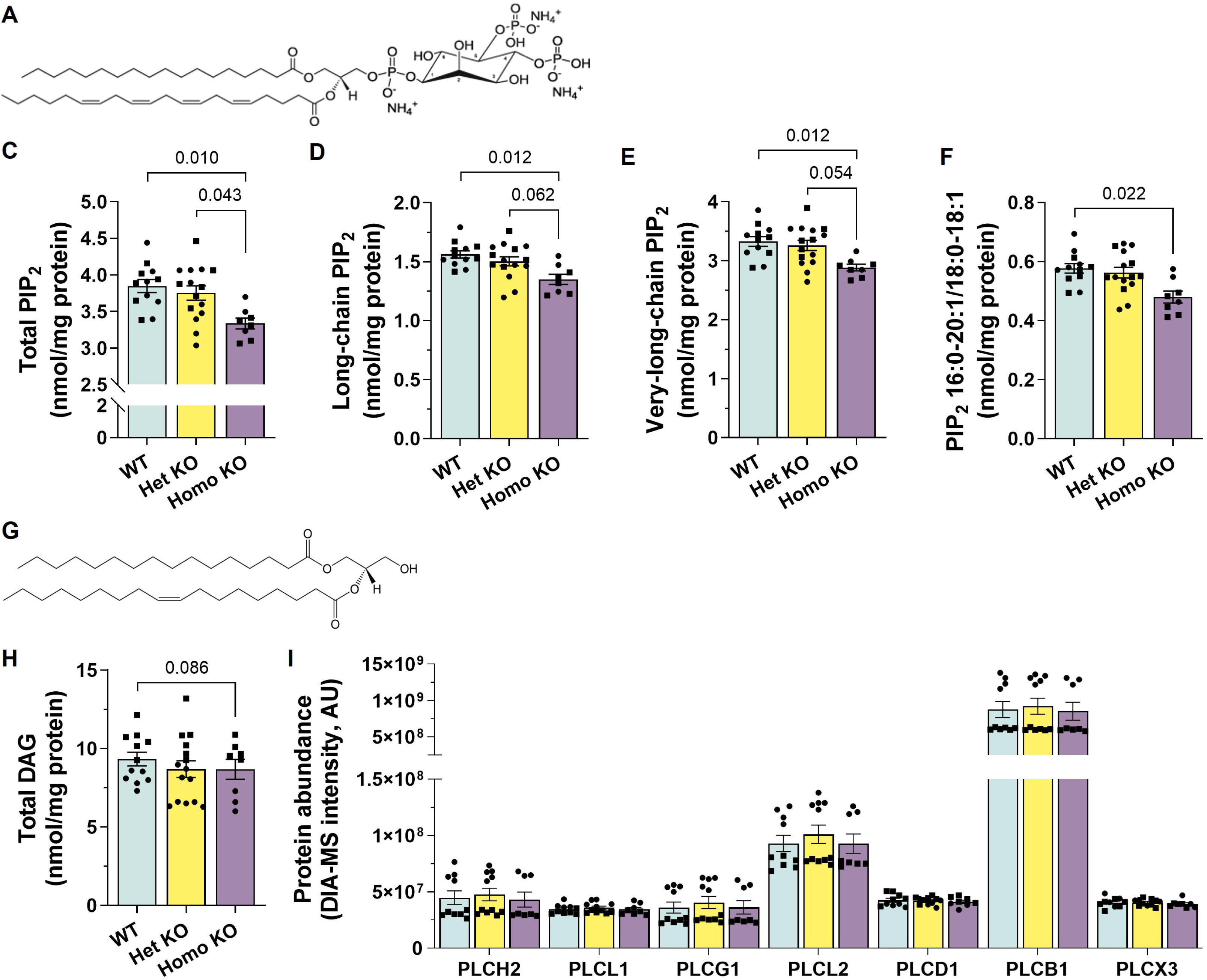
Effects of PLCγ2 deficiency on its enzymatic lipid substrate PIP_2_ and cleavage product DAG in cerebrum. (**A**) Structural schematic of phosphatidylinositol-4,5-bisphosphate (PIP_2_), highlighting the inositol-bisphosphate headgroup. (**B**-**D**) Shotgun lipidomics of cerebrum lysates from Plcg2 WT, Het KO, and Homo KO mice revealed significant reductions in total PIP_2_ (**B**). When PIP2 species were partitioned into long-chain PIP2 (**C**) and very-long-chain PIP_2_ (**D**), both subclasses were significantly reduced in Homo KO cerebrum compared with WT and Het KO littermates. (**E**) The abundant PIP2 species 16:0–20:1/18:0–18:1 was likewise decreased in Homo KO cerebrum. (**F**) Structural schematic of diacylglycerol (DAG), the canonical PLCγ2 cleavage product. (**G**) Total cerebrum DAG content was comparable across genotypes, with only a modest, non-significant reduction in Homo KO mice. Bars show mean ± SEM; individual points represent individual animals (males, squares; females, circles). FDR-adjusted p values from linear models adjusted for sex are provided in the graphs.

The female-biased negatively correlated module was enriched for oligodendrocyte differentiation, lysosomal function, regulation of macrophage migration, and signaling by interleukins, highlighting coordinated alterations in myelin, lysosomal, and immune pathways in *Plcg2*-deficient brains (Figure 5B). In contrast, the two positively correlated modules were enriched for myelin sheath and carboxy/fatty-acid metabolic processes in both sexes, with the male-biased module additionally enriched for angiogenesis and the female-biased module enriched for protein kinase binding and related signaling functions (Figure 5C–D).

### Subtle microglial transcriptional and morphological changes in Plcg2-deficient brains

Because PLCγ2 is highly enriched in microglia and functions downstream of immune receptors such as TREM2, where it modulates microglial activation and phagocytic signaling, we next sought to validate and refine the microglial signatures highlighted by our multi-omic and network analyses. NanoString Glial Panel data showed that *C1qa* and *C1qb* transcripts were decreased by ∼10% in *Plcg2* Homo KO mice compared with WT (Figure 6A–B), with *C1qc* showing a ∼10% downward trend (adj. p = 0.097). Effect sizes and expression levels for these genes were highly similar across cerebrum and brainstem and between males and females, indicating that the observed genotype effects on gene expression were not driven by a single sex or brain region.

We next attempted to validate these complement changes by qPCR in cerebrum. *C1qa* showed a clear 29% decrease in *Plcg2* Homo KO mice relative to WT that did not reach statistical significance but was directionally consistent with the NanoString data (Figure 6C). Extending this analysis to the protein level, DIA-MS proteomics revealed that C1QA and C1QB protein abundance was reduced by 20% and 14%, respectively, in *Plcg2* Homo KO cerebrum compared with WT (Figure 6D-E), with a similar 17% decrease in C1QC that trended toward but did not reach statistical significance after correcting for multiple comparisons (Figure 6F). As males and females were analyzed in separate DIA-MS runs, the consistently lower protein abundances in male samples should be interpreted cautiously, as they may reflect both biological and run-to-run effects; statistical comparisons nevertheless included both sexes with sex modeled as a covariate in a linear model. *Plcg2* Het KO mice generally showed intermediate expression levels between WT and Homo KO, typically closer to WT, consistent with a partial gene-dosage effect across the complement pathway.

Consistent with a broader shift in microglial effector programs, NanoString data showed that *Ctss* mRNA, encoding the lysosomal cysteine protease cathepsin S – a marker associated with activated and disease-associated microglia states – was reduced by 14% in *Plcg2* Homo KO mice compared with WT (Figure 6G). *Plcg2* Het KO mice again tended to show *Ctss* levels intermediate between WT and Homo KO. qPCR in cerebrum detected only a small, non-significant 4% decrease in *Ctss* (Figure 6H), again with the same direction of effect as NanoString, suggesting that the digital counting approach may be more sensitive for capturing the modest lysosomal gene changes associated with *Plcg2* deficiency.

Complementary immunofluorescence analysis demonstrated a significant reduction in Iba1⁺ area coverage by 33% within the cerebrum region of interest in *Plcg2* Homo KO mice (Figure 6I-L), while P2RY12⁺ area fraction remained unchanged across genotypes (Figure 6M-P), indicating subtle reductions in overall microglial coverage in line with the mild C1q and Ctss downregulation.

To gain a broader view of microglial activation states, we next examined NanoString pathway scores derived from curated microglial gene sets. Microglial neurodegenerative phenotype (MgND) pathway scores were markedly reduced in *Plcg2* Homo KO brains relative to WT (2.2-fold lower; Figure 6Q), suggesting reduced engagement of this neurodegeneration-associated microglial program in line with Ctss/CTSS data. Homeostatic and phagocytosis pathway scores were likewise substantially reduced (∼2.5-fold lower than WT), whereas NF-κB signaling scores were roughly doubled in Homo KO relative to WT. These pathway-level changes are consistent with the WGCNA module enriched for immune and lysosomal genes negatively associated with *Plcg2* genotype and, together with the concordant shifts in microglia-enriched transcripts and proteins and in Iba1 and P2RY12 immunostaining.

### Coordinated lipid and amino acid catabolism in Plcg2-deficient brains revealed by acylcarnitine metabolomics

Within the NanoString dataset, two of the 11 upregulated DEGs were linked to oxidative and peroxisomal lipid pathways. Absolute NanoString quantification showed that Phyh mRNA was increased by 6% in Plcg2 Homo KO cerebrum compared with WT (Figure 7A). Fth1 transcript was increased by 7% in Plcg2 Homo KO cerebrum (Figure 7B). Effect sizes for Phyh and Fth1 were similar in males and females, and absolute expression levels were higher in brainstem than cerebrum.

In the WGCNA analysis, modules positively associated with Plcg2 Homo KO status were enriched for Gene Ontology terms, including “fatty-acid metabolic process” and “carboxylic-acid metabolic process” (Figure 5C–D). STRING and Enrichr analyses of differentially abundant proteins identified KEGG pathways related to mitochondrial lipid and amino-acid catabolism (Figure 7C–D). DIA-MS proteomics showed that CPT1A and HADHA protein abundance was increased by approximately 10% in Plcg2 Homo KO cerebrum relative to WT (Figure 7E–F). ACADSB was elevated by 10%, and ALDH9A1 by 14%, in Plcg2 Homo KO cerebrum (Figure 7G, Figure S3A).

Gene Ontology cellular-component analysis identified the mitochondrial matrix as the top enriched term for this protein set (Figure 7D). Within this matrix-localized group, ETFB abundance was increased by 8% in Plcg2 Homo KO brains (Figure 7H), and additional matrix enzymes including ACSS1, RIDA, and HMGCS1 showed increases of 13–22% (Figure S3B–D).

Based on these findings, we performed LC/MS-based targeted acylcarnitine profiling using isotope-labeled internal standards for each species in an independent cohort of 3-month-old PBS-treated Plcg2 WT, Het KO, and Homo KO mice (Figure 7I–J). A metadata heatmap and hierarchical clustering of MetaboAnalyst annotations showed that Homo KO brains clustered together irrespective of sex, whereas WT and Het KO samples grouped primarily by sex (Figure 7I). PCA of acylcarnitine profiles with genotype as the grouping factor revealed substantial overlap between WT and Het KO samples and a separate cluster of Homo KO samples (Figure 7J). PERMANOVA on the acylcarnitine distance matrix indicated a significant main effect of genotype.

To identify individual acylcarnitines associated with Plcg2 genotype, we applied a linear model with sex as a covariate in MetaboAnalyst. Among 27 quantified species (C0 and C2–C20), three showed significant genotype effects after FDR correction: 3-hydroxyhexanoylcarnitine (C6-OH), 2-octenoylcarnitine (C8:1), and L-glutarylcarnitine (C5-DC) (Figure 7K–L). C6-OH and C8:1 were increased by 36% and 70%, respectively, in Homo KO brains relative to WT, and C6-OH was also increased relative to Het KO (Figure 7K–L).

An exploratory Spearman rank correlation analysis in MetaboAnalyst showed that six acylcarnitine species (C18:2, C8:1, C14:2, C12:1, C5-DC, and C6-OH) displayed absolute Spearman ρ values greater than 0.5 with Plcg2 genotype (Figure S3E). Short-, medium-, and long-chain acylcarnitines with strong genotype correlations showed increased levels in Plcg2 Homo KO brains, whereas free carnitine (C0) did not differ significantly across genotypes.

### Selective PIP_2_ depletion in Plcg2-deficient brains despite low bulk PLCγ2 abundance

Given that PLCγ2 directly hydrolyzes phosphatidylinositol-4,5-bisphosphate (PIP_2_) to yield diacylglycerol (DAG) and inositol trisphosphate, we next examined cerebrum PIP_2_ and DAG content in Plcg2-deficient brains. Total levels of PIP_2_, the canonical substrate of PLCγ2, were reduced by ∼13% in Plcg2 Homo KO brains compared with WT (Figure 8A-B), with similar significant decreases across long-chain and very-long-chain PIP_2_ suclasses (Figure 8C-D). Five of the differentially abundant lipids were PIP_2_ species, including the second most abundant molecular species, PIP_2_ 16:0–20:1/18:0–18:1, which showed a concordant 17% decrease in Homo KO brains (Figure 8E). Similarly, the most abundant species, PIP_2_ 18:0–20:4, tended to decrease by ∼13% but did not remain significant after correction for multiple comparisons (Figure S4A).

Despite this selective depletion of brain PIP_2_ pools, total levels of phosphatidylinositol-4-phosphate (PIP_1_) and phosphatidylinositol-3,4,5-trisphosphate (PIP_3_) were unchanged across genotypes (Figure S4B–C), indicating that *Plcg2* deficiency does not globally disrupt phosphoinositide abundance. Finally, total diacylglycerol (DAG) levels only modestly trended downward in Homo KO brains relative to WT (7% reduction; Figure 8F-G). Trying to get more granularity we broke into saturation-based DAG subclasses (saturated, monounsaturated, polyunsaturated), but none were significantly altered (Figure S4D-F).

Consistent with the notion that PLCγ2 is a low-abundance signaling node in bulk brain, DIA-MS profiling of PLC-family proteins in whole cerebrum revealed robust expression of only seven out of 18 PLC-family members, comprising 16 mammalian PLC isozymes and two PLC-like proteins (PLCL1/2). PLCB1 dominated total PLC abundance and was the only detectable PLCβ isozyme, followed by PLCH2 as the only detectable η-family isozyme, PLCG1 as the only detectable γ-family isozyme, and PLCD1 as the only detectable δ-family isozyme (Figure 8H). Notably, both PLC-like proteins PLCL1 and PLCL2 were detected, with PLCL2 emerging as the second most abundant PLC-family protein in whole cerebrum (Figure 8H). In contrast, PLCγ2 protein and 10 additional PLC isozymes were below the limit of detection in these whole-cerebrum lysates, indicating that the observed PIP_2_ depletion occurs in a context where PLCγ2 contributes only a minor fraction of total PLC protein in bulk brain. Notably, none of the robustly detected PLC-family members are selectively enriched in microglia based on single-cell and single-nucleus transcriptomic datasets, in contrast to PLCγ2, which is predominantly expressed by microglia in the brain and functions as a key signaling node downstream of TREM2 and other immune receptors.

### Plcg2 deficiency reveals an unexpected role in central myelin homeostasis

As evidenced by the lipidomics (Figure 4C) results, the most prominent myelin-linked signal was a coordinated reduction in HexCer species, with roughly half of the differentially abundant lipids belonging to this class. Because most of brain HexCer corresponds to galactosylceramides (galactocerebrosides) that predominantly localized to compact CNS myelin, even modest changes in this class are expected to primarily reflect altered myelin lipid content. Total HexCer was reduced by ∼15% in Plcg2 Homo KO cerebrum relative to WT and Het KO mice (Figure 9A-B), and this effect was primarily driven by losses in the non-hydroxy HexCer subclass by 23% (Figure 9C), whereas hydroxy HexCer species showed a downward trend (-8%) that did not reach statistical significance. The most abundant galactosylceramide species, HexCer N24:1, was decreased by ∼20% in Homo KO brains (Figure 9D), reinforcing that PLCγ2 deficiency selectively targets the galactosylceramide-rich myelin lipid pool.

**Figure 9.**
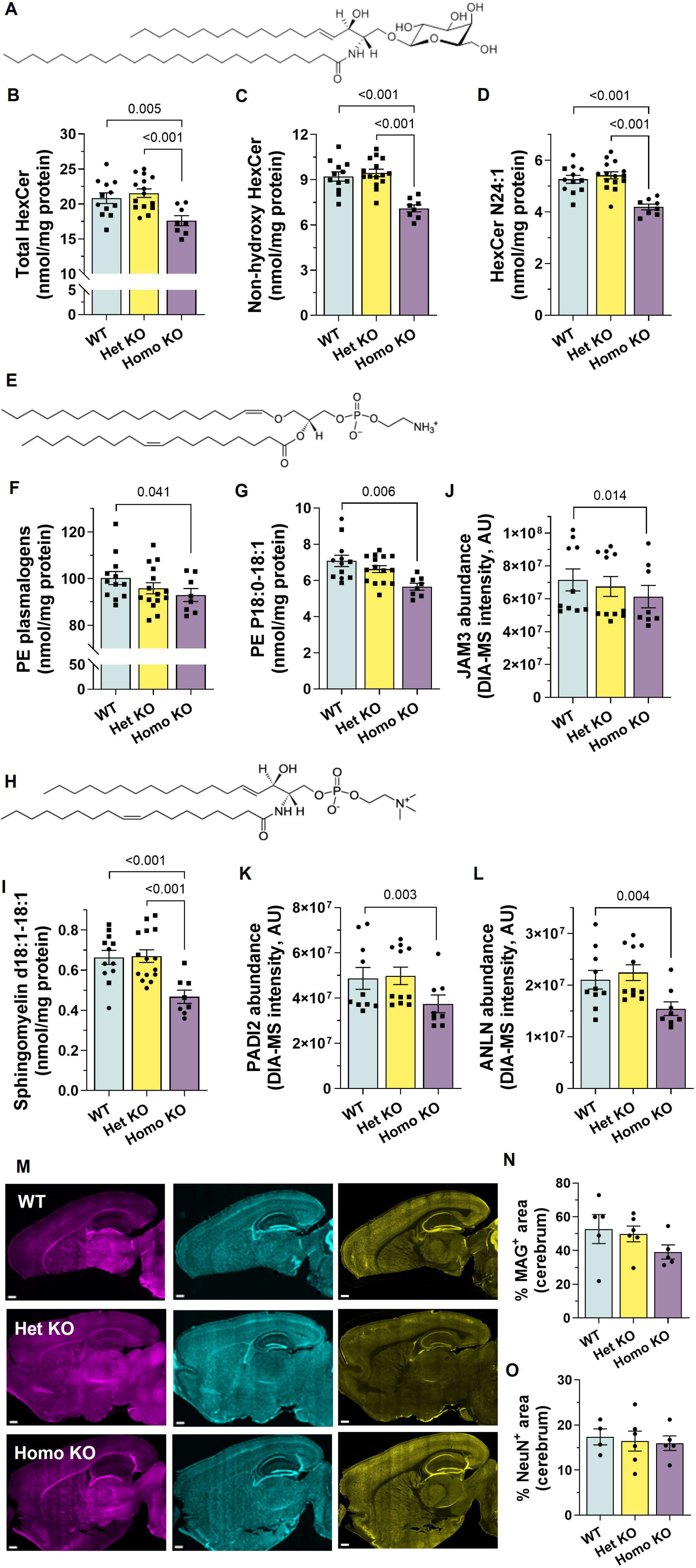
Myelin-enriched lipids and myelin-associated proteins in Plcg2-deficient brains. (**B**-**D**) Shotgun lipidomics was used to quantify hexosylceramides (HexCer), primarily composed of glucocerebroside in cerebrum, including total HexCer, non-hydroxy HexCer, and the abundant HexCer N24:1 species, in Plcg2 WT, Het KO, and Homo KO mice. (**F**–**G**) Phosphatidylinositol-4,5-bisphosphate (PIP_2_) levels were measured as total PIP_2_ and a representative molecular species, PIP_2_ 16:0–20:1/18:0–18:1. (**H**–**J**, **L**) Additional myelin-enriched lipid subclasses, including ethanolamine plasmalogens and the plasmalogen species PE P18:0–18:1, as well as the sphingomyelin species SM d18:1–18:1, were quantified in the same samples. (**I**-**K**) DIA-MS proteomics was used to measure abundance of myelin- and oligodendrocyte-enriched proteins, including JAM3, PADI2, and ANLN, in cerebrum. (**M**-**O**) Immunofluorescence for myelin-associated glycoprotein (MAG), NeuN, and GFAP was used to assess myelin, neuronal, and astrocytic area fractions in sagittal brain sections. Data are presented as bar plots, with each point representing an individual animal (males, squares; females, circles); statistical tests and multiple-comparison corrections applied to lipid, proteomic, and immunohistochemical measurements are described in the Methods. Ordinary one-way ANOVA with Tukey’s post hoc tests was used for panels L and P.

Subclass analysis reinforced this pattern of myelin-linked vulnerability. In addition to non-hydroxy HexCer and the long- and very-long-chain PIP_2_ subclasses, ethanolamine plasmalogens emerged as the only other lipid subclass that met significance, and this class is likewise highly enriched in myelin membranes. The phosphatidylethanolamine plasmalogen subclass was significantly decreased in Plcg2 Homo KO cerebrum (Figure 9E-F), exemplified by reduced levels of the abundant PE plasmalogen species PE P18:0–18:1 (Figure 9G). Because ethanolamine plasmalogens (a.k.a., plasmenylethanolamine) are highly enriched in central myelin, their coordinated decline alongside HexCer further supports subtle erosion of myelin membrane integrity rather than a global lipid deficit. Finally, we observed a significant decrease in a sphingomyelin species, SM d18:1-18:1 (Figure 9H-I), mirrored by trending decreases in SM d18:1-24:0 and d18:1-24:1.

Consistent with the lipidomics signature, DIA-MS proteomics revealed that several proteins with strong enrichment in myelin or oligodendrocyte compartments were differentially abundant in Plcg2 Homo KO cerebrum. Among these, JAM3 (junctional adhesion molecule 4), PADI2 (peptidyl arginine deiminase 2), and ANLN (anillin) were significantly decreased, each showing modest but consistent shifts in abundance that mirror the direction of the myelin lipid changes (Figure 9J–L). These proteins have been implicated in paranodal junction organization, myelin protein citrullination, and cytoskeletal dynamics, respectively [39, 59, 60].

To determine whether these biochemical changes translate into detectable alterations in myelin architecture, we quantified myelin-associated glycoprotein (MAG) immunoreactivity in sagittal brain sections (Figure 9M). MAG-positive area tended to be lower in *Plcg2* Homo KO mice compared with WT littermates (Figure 9N), aligning with the selective loss of myelin-enriched lipid species. In contrast, NeuN-positive neuronal nuclei (Figure 9O) and GFAP-positive astrocytes were unchanged across genotypes.

## Discussion

### Developmental and vascular roles of PLCγ2

Our data reinforce that PLCγ2 is essential for normal developmental survival and vascular patterning. Plcg2 Het × Het intercrosses yielded far fewer Homo KO weanlings than predicted by Mendelian ratios, with genotype distributions indicating substantial pre- and early postnatal lethality and implying that roughly half of Plcg2-deficient embryos die in utero and many of the remainder succumb between P0 and P5. Survival was sexually dimorphic, with female Homo KO animals exhibiting higher mortality than males, echoing the female bias in multiple autoimmune conditions [61, 62] and the emerging view that immune regulatory pathways intersect with sex-dependent longevity biology [61].

In surviving young adult Homo KO mice, gross examination revealed prominent intestinal vascular abnormalities characterized by a dense meshwork of thin, highly arborized blood-filled vessels that persisted after saline perfusion and appeared more marked in females. These intestinal findings are concordant with prior Plcg2-null and Plcg2^al/al^ models, in which disrupted blood-lymphatic separation leads to blood-filled lymphatic vessels, gastrointestinal hemorrhage, and fatal hemodynamic compromise [22, 63], and the convergence of these phenotypes across independent alleles—including the MODEL-AD CRISPR/Cas9-engineered knockout used here—supports a model in which PLCγ2 acts as a non-redundant regulator of blood–lymphatic vascular partitioning.

Despite the high burden of early mortality and intestinal vascular disruption, the subset of Plcg2 Homo KO mice that survived to weaning reached three months of age with normal body weight and hemicerebrum mass, suggesting either partial vascular compensation or selective survival of individuals with less severe developmental defects. However, non-fasted blood glucose was modestly but reproducibly reduced in Homo KO mice, with intermediate values in Het KO animals, indicating that altered systemic metabolic homeostasis is already detectable in young adulthood. This shift could reflect direct contributions of PLCγ2 signaling to peripheral glucose regulation or, alternatively, chronic vascular and immunologic stress that secondarily influences endocrine and metabolic set points; more detailed metabolic and endocrine profiling will be required to distinguish these possibilities and to determine how they intersect with the AD-relevant PLCG2 variant spectrum.

### Systemic immune remodeling and impaired immune tolerance

Beyond developmental and vascular phenotypes, Plcg2 deficiency produced a characteristic pattern of systemic immune remodeling, marked by splenomegaly, myeloid-skewed spleen composition, enrichment of age-associated B-cell (ABC-like) subsets, and parallel reductions in CD4 and CD8 regulatory T-cell populations. These anatomical changes are consistent with high *Plcg2* expression across B-cell developmental stages in spleen and in blood myeloid progenitor and effector lineages in the GEXC hematopoiesis expression map, as well as with prior reports of immune abnormalities in Plcg2-deficient mice [22, 64, 65].

Splenomegaly in Plcg2 Homo KO animals aligns with earlier work showing increased spleen cellularity in Plcg2 knockout mice [22], but our data indicate that this enlargement does not reflect expansion of the mature B-cell compartment, which instead is reduced in proportion and absolute number. The decrease in CD19⁺CD11b⁻ B cells is consistent with impaired BCR-dependent survival signaling and reduced B cell maturation described in Plcg2-deficient models, whereas CD19⁻CD11b⁺ cells, a population enriched for myeloid/innate lineages (e.g., monocytes/macrophages and granulocytes), are increased in both percentage and number, supporting the idea that splenomegaly in this context reflects accumulation of myeloid cells with defective migration due to loss of Plcg2 [37], rather than generalized lymphoid expansion. Given that PLCγ2 is required for chemokine- and integrin-coupled signaling in multiple myeloid lineages, these shifts raise the possibility that defective migration or tissue egress contributes to myeloid buildup in spleen in the absence of PLCγ2. Collectively, these changes indicate that PLCγ2 deficiency does not simply remodel spleen size, but rebalances peripheral immunity away from conventional B cells and toward myeloid/innate effectors, reducing the pool of cells that normally support humoral homeostasis. In this context, the concomitant expansion of pro-inflammatory ABC-like CD11c⁺T-bet⁺ B cells and depletion of CD4 and CD8 regulatory T-cell subsets removes key tolerance-maintaining lymphocytes while enriching effector populations that drive chronic inflammation, providing a cellular basis for compromised systemic immune tolerance in Plcg2-deficient mice.

The prominent increase in ABC-like B cells in spleen and brain-draining cervical lymph nodes is notable because these cells represent a pro-inflammatory B-cell subset that accumulates with aging and autoimmunity and is typically associated with chronic antigenic and innate stimulation [26, 66, 67]. At the same time, FoxP3⁺ CD4 Tregs and CD8 Tregs are significantly reduced in both spleen and cervical lymph nodes, a pattern that mirrors enhancement of aging- and autoimmune-like phenotypes with PLCγ2 deficiency and aligns with reports that PLCγ2 hypermorphic variants can increase Treg numbers [68]. Because the hypermorphic variant and the complete knockout represent opposite extremes of PLCγ2 signaling, the Treg decrease we observe in Homo KO mice is conceptually consistent with a model in which PLCγ2 activity supports Treg homeostasis. Because regulatory T cells serve as key brakes on age- and autoimmunity-associated immune activation, their depletion is expected to promote a chronic low-grade inflammatory milieu characterized by elevated interferon γ [69], such a cytokine environment is well-suited to sustain T-bet⁺ ABC-like populations [70].

Therefore, the depletion of T regulatory cells may provide a permissive environment for production of ABC-like cells. However, PLCγ2 is a key signaling effector downstream of the BCR and ABCs are generally thought to depend on chronic BCR signaling for their generation and maintenance [71], so it is not entirely clear how ABC-like cells increase in PLCγ2-deficient mice. With respect to loss of T regulatory subsets, this may be due to altered T cell development found with PLCγ2 deficiency [23] or secondary to myeloid cell dysfunction with due to PLCγ2 deficiency.

The fact that both ABC enrichment and Treg depletion are evident in cervical lymph nodes is particularly relevant to neuroimmunology, as these nodes drain meningeal and glymphatic outflow and thus integrate immune signals at the brain-border interface. Together, these findings support a model in which PLCγ2 deficiency drives a coordinated shift toward pro-inflammatory, ABC-rich B-cell states and diminished regulatory T-cell control in both systemic and brain-draining lymphoid tissues, a combination that is predicted to compromise immune tolerance and sensitize to autoimmune or neuroinflammatory pathology over time. This is especially relevant given that *PLCG2* variants are implicated in both autoinflammatory disease and human longevity [4], linking PLCγ2 to the broader interface of immune regulation, autoimmunity, neurodegenerative risk, and lifespan.

### Neuroimmune context: PLCγ2 in microglia and brain-border immune niches

Public single-cell RNA-seq datasets converge on the view that PLCγ2 is a low-abundance but selectively expressed signaling enzyme in brain-resident and border-associated immune compartments. In the Aging Mouse Brain Single Cell Portal, *Plcg2* expression is highest in microglia and brain macrophages, with lower levels in endothelial cells and oligodendrocyte precursor cells and minimal to no expression in neurons, astrocytes, and mature oligodendrocytes [30, 72–74]. Virtually identical expression patterns are observed in the DropViz adult mouse brain atlas, where the top ten *Plcg2*-expressing clusters are classified as microglia/macrophages and endothelial cells, with the strongest expression in microglia/macrophage clusters marked by Tmem119-Mrc1 and C1qb [73].

Because *Plcg2* expression in these datasets is concentrated in both myeloid and vascular compartments, we next examined single-cell atlases that resolve endothelial and border-associated immune populations. The Betsholtz lab vascular atlas shows highest *Plcg2* expression in microglia, followed by arterial endothelial cells, highlighting putative roles at the blood-brain and perivascular interfaces [72]. In the Brain Immune Atlas aggregate of CD45⁺ cells from brain and meningeal regions, *Plcg2* transcripts are detected in 18% of immune cells across parenchymal microglia and multiple border-associated macrophage (BAM) subsets, with choroid plexus surface and dural BAMs contributing many *Plcg2*⁺ cells and a small B-cell cluster displaying the highest per-cell *Plcg2* expression in the dataset [74]. In this atlas, the distribution of *Plcg2* more closely parallels the ITAM-associated kinase *Syk* than Trem2, *Bcr*, or *Tlr4*, supporting a model in which PLCγ2 acts as a shared signaling hub downstream of Syk-coupled receptors in microglia, BAMs, meningeal B cells, and arterial endothelial cells.

Complementary single-cell analyses further refine this picture by identifying PLCγ2-high age-associated B-cell–like clusters at CNS borders. An unpublished mouse atlas of CNS-resident B cells reveals enrichment of *Plcg2* in CD11c⁺T-bet⁺ ABC-like clusters in meningeal and brain-associated lymphoid tissues, with analogous PLCG2-high ABC-like populations in human post-mortem brain B-cell datasets. Taken together, these datasets reinforce that microglia are the predominant *Plcg2*-expressing cell type within the brain parenchyma, while also identifying BAMs, ABC-like B cells at CNS borders, and arterial endothelial cells as *Plcg2*-enriched populations that are well positioned to mediate PLCG2-modulated neuroimmune and vascular signaling. Across all datasets, *Plcg2* transcript counts remain relatively low even in these populations, consistent with PLCγ2 being a low-abundance yet selectively expressed enzyme under physiological, non-inflammatory conditions.

In concordance with this atlas-based context, our multi-omic profiling indicates that PLCγ2 loss subtly but consistently reshapes microglial effector programs in young adult mice. Across RNA and protein levels, we detected modest reductions in microglia-enriched complement components (C1q subunits) and the lysosomal protease Ctss, together with decreased scores for neurodegeneration-associated and phagocytic microglial gene sets and a relative increase in NF-κB signaling pathway scores. These coordinated changes support the conclusion that PLCγ2 deficiency in young adult mice is associated with mild but systematic alterations in microglial complement, lysosomal, and signaling programs, dampening complement- and lysosome-linked microglial states while shifting signaling balance toward NF-κB-associated responses even in the absence of overt neurodegeneration. This configuration is consistent with prior work showing that loss of PLCγ2 can increase NF-κB signaling while reducing activation of the interferon regulatory factor 3 (IRF3) pathway in PLCγ2-deficient macrophages [75]. Because these measurements were obtained in young adult mice with minimal baseline neuroinflammation, even relatively small decrements in complement and lysosomal gene expression could become functionally more consequential under inflammatory or amyloid-driven conditions.

Morphological and flow-cytometric data point to a parallel, subtle shift in microglial phenotype. In PLCγ2-deficient mice, we observed a slight decrease in Iba1⁺ brain myeloid coverage by immunostaining, no change in P2RY12⁺ microglial area, and a small increase in CD11c mean fluorescence intensity in microglia by flow cytometry. Taken together, these changes are consistent with a modest phenotypic shift in microglia that preserves core homeostatic features while altering others. CD11c⁺ microglia are known to increase in chronic inflammatory and neurodegenerative conditions as well as during development [76–83], and CD11c⁺ microglia have been implicated in myelin development and repair [79, 84], raising the possibility that these CD11c-high microglia may be responding to—or even contributing to—the myelin lipid and protein changes we observed by lipidomics and proteomics.

At the network level, WGCNA-derived co-expression modules align with these microglial findings and extend them to broader glial and vascular programs. Plcg2-sensitive modules are enriched for microglial/immune and lysosomal pathways, myelin-associated transcripts, fatty-acid metabolism, and vascular signaling, suggesting that PLCγ2 deficiency remodels coordinated gene networks rather than isolated genes. Female-biased positively correlated modules show enrichment for protein kinase binding and related signaling terms, consistent with PLCγ2’s regulation by upstream tyrosine kinases and its role in generating DAG to activate PKC and other kinase-dependent pathways, whereas male-biased positively correlated modules are enriched for angiogenesis-related pathways, in line with known roles for PLCγ2 in vascular and lymphatic development and blood–lymphatic separation. Together, these WGCNA results provide a framework that links microglial complement and activation signatures, myelin-associated transcripts, fatty-acid oxidation pathways, and vascular signaling into integrated PLCγ2-sensitive networks, and they motivated our targeted validation of microglial, myelin, and oxidative phenotypes in Plcg2-deficient brains.

Viewed alongside our in vivo data showing ABC enrichment and Treg loss in cervical lymph nodes, these microglial and network-level changes support a coherent picture in which PLCγ2 deficiency biases both central and peripheral compartments toward innate-like, inflammation-associated immune states with impaired immune tolerance. In this model, microglia, BAMs, arterial endothelial cells, and ABC-like B cells at CNS borders constitute a distributed PLCγ2-dependent signaling axis that integrates neuroimmune and vascular cues and is poised to influence how the brain responds to amyloid, tau, and other neurodegenerative stressors over the lifespan.

### Selective myelin vulnerability and PIP_2_ loss in Plcg2-deficient brain

Multiple lines of evidence from our study converge on myelin as a key node of PLCγ2-mediated brain homeostasis. First, the strongest lipidomic signal of Plcg2 deficiency was a coordinated reduction in myelin-enriched lipid classes: total HexCer was reduced, driven by non-hydroxy HexCer species that correspond predominantly to galactosylceramides, together with significant decreases in ethanolamine plasmalogens. These changes are unlikely to reflect a global lipid deficit, because other major classes were largely preserved, and they map instead onto lipid subclasses that are specific to compact CNS myelin. Second, proteomics and histology support a subtle structural correlate: several myelin- and paranode-associated proteins (e.g., JAM4, PADI2, ANLN) were decreased, and MAG-positive white matter area tended to be lower in Plcg2 Homo KO brains, without accompanying loss of NeuN or increase in GFAP, providing an additional readout that converges with the HexCer and plasmalogen changes on lipid components of myelin membranes and further linking PLCγ2 deficiency to altered myelin structure and turnover at the protein level. Third, transcriptomic profiling revealed increased expression of oligodendrocyte/myelin-related genes such as Mog, Cnp, Plp1, Bcas1, Fth1, and Phyh, suggesting that oligodendrocytes mount a compensatory transcriptional response to maintain myelin integrity as lipid content declines.

These converging biochemical, proteomic, histologic, and transcriptional data strongly argue that PLCγ2 deficiency produces a mild but coherent erosion of myelin membranes that is actively countered at the RNA level, rather than a nonspecific injury to all brain cell types. Our findings thus position PLCγ2 as an unexpected modulator of white matter homeostasis, supporting a model of subtly compromised myelin integrity in PLCγ2-deficient brains and highlighting potential relevance for AD-associated white matter vulnerability. Within this myelin-centered framework, the cerebrum PIP2 phenotype takes on a different meaning. Total PIP2 and virtually all PIP2 species were reduced (significantly or borderline) in Plcg2 Homo KO cerebrum, a result that appears counterintuitive if one assumes that PLCγ2 loss should lead to substrate accumulation. This reduction occurred without detectable compensatory changes in other PLC isoforms at the RNA or protein level, and DAG and other phosphoinositide subclasses were largely preserved, pointing to a selective impairment of cerebrum PIP2 homeostasis in the absence of PLCγ2 rather than a broad collapse of PLC-coupled phosphoinositide and DAG signaling.

Within this myelin-centered framework, the PIP_2_ phenotype takes on a different meaning. Total PIP_2_ and virtually all PIP_2_ species were reduced (significantly or borderline) in Plcg2 Homo KO cerebrum, a result that appears paradoxical if one assumes that PLCγ2 loss should lead to substrate accumulation. However, two observations support the alternative interpretation that bulk PIP_2_ reductions primarily reflect loss of a PIP_2_-enriched myelin compartment rather than direct depletion of PIP_2_ by PLCγ2: (i) in our proteomic dataset, PLCγ2 protein itself was not detectable in cerebrum, despite sensitive DIA-MS and broad coverage of other PLC isoforms, indicating that PLCγ2 is a very low-abundance enzyme in the resting adult brain; and (ii) the PIP_2_ species that decline in Plcg2-null brains are those with long and very-long-chain acyl compositions typical of myelin-enriched phosphoinositides.

These observations raise the possibility that PLCγ2 contributes disproportionately to maintaining brain PIP2 pools, or that PIP2 depletion is secondary to other PLCγ2-dependent processes. In light of the coordinated changes in myelin-enriched lipids described above, a parsimonious explanation is that loss of PLCγ2 perturbs oligodendrocyte and myelin signaling pathways that normally sustain myelin-associated PIP2, coupling altered PIP2 homeostasis to emerging deficits in central myelin. Consistent with this idea, the PIP2 species that decline in Plcg2-null brains are enriched for long and very-long-chain acyl compositions typical of myelin-associated phosphoinositides. Published work further supports the notion that PIP2 is enriched in myelin membranes and is functionally linked to myelin biology: multiple studies using myelin-enriched fractions or white matter-focused lipidomics have reported substantial concentrations of phosphoinositides, including PIP2, within CNS myelin, and have implicated PIP2 in regulating myelin protein trafficking, cytoskeletal coupling, and paranodal organization [39–41]. In demyelinating and dysmyelinating models, phosphoinositide distributions within white matter tracts are altered [85–87] consistent with a tight coupling between myelin integrity and local PIP_2_ pools. In light of these data, the simplest explanation for our observations is that PIP_2_ species behave as myelin-enriched phosphoinositides, and that their decline in Plcg2 Homo KO cerebrum is largely a consequence of selective myelin lipid loss rather than a primary effect of PLCγ2 on PIP_2_ turnover.

Although our data favor a myelin-enriched interpretation of the PIP_2_ phenotype at the tissue level, single-cell atlases and our own data mining indicate that Plcg2 is highly expressed in microglia and brain-associated macrophages, and that a subset of “immune oligodendrocytes” also expresses Plcg2 at levels approaching those in microglia. These observations leave open the possibility of more direct, microdomain-restricted roles for PLCγ2 in specific PIP_2_ pools, for example at microglial processes that surveil and remodel myelin. More broadly, we propose that PLCγ2 may be a central regulator of white-matter-associated microglia that normally clear physiological myelin debris and support remyelination, such that loss of PLCγ2 activity impairs microglia-oligodendrocyte crosstalk and myelin turnover, leading over time to the subtle but coordinated erosion of myelin lipids and PIP_2_-rich myelin membranes observed here. Future studies using myelin fractionation, cell type-specific *Plcg2* deletion in microglia and oligodendrocyte lineages, and *in vivo* models of demyelination and remyelination will be needed to formally test this hypothesis and to determine whether graded PLCγ2 loss-of-function states recapitulate the myelin and PIP2 phenotypes we observe in complete knockouts.

Taken together with our oxidative metabolism findings, these myelin and PIP2 data indicate that PLCγ2 deficiency installs a coordinated program in the mouse brain characterized by impaired PIP2 homeostasis, selective loss of myelin-enriched lipid and protein components, and increased reliance on mitochondrial and peroxisomal oxidative pathways. This extends PLCγ2’s role beyond microglial signaling into broader control of myelin integrity and metabolic resilience and provides a mechanistic framework for how PLCγ2 loss-of-function variants might heighten vulnerability to Alzheimer’s disease and aging-related brain stress.

### Brain oxidative metabolism in Plcg2 deficiency

Our multiomic analyses revealed that PLCG2 loss engages a coordinated oxidative metabolism program in the young adult brain, with convergent evidence from transcriptomics, proteomics, and acylcarnitine profiling. Within the NanoString dataset, modest but significant upregulation of Phyh and Fth1 in Plcg2 Homo KO cerebrum, with higher absolute levels in brainstem, is consistent with enhanced peroxisomal oxidation of branched-chain fatty acids and increased iron-storage capacity in myelin-rich white matter tracts. In this context, upregulation of *Fth1* and *Phyh* is particularly notable: ferritin heavy chain–mediated iron delivery and storage are critical for oligodendrocyte iron metabolism and myelin lipid synthesis, helping to constrain iron-driven lipid peroxidation and ferroptotic death in white matter [88, 89], whereas the *Phyh*-encoded peroxisomal enzyme phytanoyl-CoA 2-hydroxylase supports α-oxidation of 3-methyl branched-chain fatty acids such as phytanic acid, a peroxisomal processing step that is particularly important in lipid-rich, myelinated tissues [90].

These transcript-level changes dovetail with our WGCNA results, in which modules positively associated with Plcg2 Homo KO status were enriched for “fatty-acid metabolic process” and “carboxylic-acid metabolic process,” pointing to broader remodeling of mitochondrial catabolism rather than isolated shifts in single genes. STRING and Enrichr analyses of differentially abundant proteins identified a tightly connected network centered on mitochondrial lipid and amino-acid catabolism, localized predominantly to the mitochondrial matrix and including CPT1A, HADHA, ACADSB, ALDH9A1, ETFB, ACSS1, RIDA, and HMGCS1. CPT1A catalyzes the rate-limiting formation of long-chain acylcarnitines at the outer mitochondrial membrane, thereby governing entry of long-chain fatty acids into the β-oxidation pathway [91], whereas HADHA is the catalytic subunit of the mitochondrial trifunctional protein that performs downstream dehydrogenation and hydration steps for long-chain acyl-CoAs [92]. ACADSB acts as a short/branched-chain acyl-CoA dehydrogenase for BCAA-derived and other branched substrates, and ALDH9A1 supports oxidation of amino-derived aldehydes such as 4-trimethylaminobutyraldehyde and γ-aminobutyraldehyde, contributing to carnitine biosynthesis and GABA production. The composition and mitochondrial-matrix enrichment of this module support the view that Plcg2 deficiency augments oxidative handling of both lipid-derived and amino-acid–derived substrates in the brain.

Targeted acylcarnitine metabolomics provides functional support for this integrated oxidative signature. Homo KO brains exhibit a dominant, sex-independent acylcarnitine profile, in contrast to WT and Het KO samples that segregate primarily by sex, and multiple short-, medium-, and long-chain acylcarnitines track closely with Plcg2 dosage in both linear models and correlation analyses. The strong association of dicarboxy-acylcarnitines such as glutarylcarnitine (C5-DC), generated during lysine/tryptophan-linked amino-acid oxidation, suggests that amino-acid–derived acylcarnitines are integral components of this PLCγ2-sensitive oxidative program.

Together, these findings support a model in which Plcg2 loss activates an integrated peroxisomal–mitochondrial oxidative module that helps dispose of altered lipid species and amino-acid-derived substrates in the brain. In the context of our lipidomics data, it is tempting to speculate that this oxidative reprogramming partly reflects increased turnover of myelin-associated lipids and phosphoinositide-derived fatty acids, with Phyh-driven peroxisomal α-oxidation and Fth1-mediated iron buffering helping to constrain iron-driven lipid peroxidation and ferroptotic stress in vulnerable white matter regions. Although we did not detect overt neurodegeneration or gliosis at three months, the coordinated engagement of peroxisomal–mitochondrial oxidative pathways, together with subtle myelin lipid loss, may shape how Plcg2-deficient brains respond to aging, inflammatory, and AD-related stressors that challenge white matter integrity and oxidative resilience.

### Implications for PLCG2 variants in neurodegeneration and aging

Human genetic studies have identified a nonsynonymous *PLCG2* variant (rs72824905-G) that confers reduced risk of AD, dementia with Lewy bodies, and frontotemporal dementia while increasing the likelihood of longevity, highlighting PLCG2 as a candidate therapeutic target [4]. Our data underscore that complete loss of Plcg2 function produces deleterious developmental, vascular, and immune phenotypes, suggesting that protective human variants are unlikely to act as simple loss-of-function alleles. Instead, they more plausibly exert partial, context- and cell-type-specific modulation of PLCγ2 signaling.

Within this framework, the ABC expansion, Treg depletion, and early microglial and myelin-associated changes observed in Plcg2-deficient mice can be viewed as exaggerated manifestations of pathways that are more subtly tuned by protective *PLCG2* alleles in humans. On the systemic side, Plcg2 loss pushes the immune system toward a pro-inflammatory, ABC-rich and Treg-poor state that is expected to weaken peripheral tolerance and favor chronic low-grade inflammation, particularly in females, who already show higher susceptibility to autoimmune disease and exhibited greater mortality in our colony. On the neuroimmune side, PLCγ2 sits at a shared signaling hub in microglia, border-associated macrophages, arterial endothelial cells, and ABC-like B cells, and its absence is associated with coordinated alterations in microglial complement and lysosomal programs, selective myelin-associated lipid and protein loss, impaired PIP2 homeostasis, and increased reliance on mitochondrial and peroxisomal oxidative pathways. These pleiotropic roles across vascular, systemic immune, and neuroimmune compartments intersect with key mechanisms of aging and neurodegeneration.

Altogether, our findings support a model in which PLCγ2 acts as a multi-compartmental integrator of immune, vascular, and metabolic signals that collectively shape neurodegenerative risk. In this model, protective *PLCG2* variants may preserve or enhance adaptive microglial and border-associated responses to amyloid and other insults, stabilize myelin and vascular integrity, and fine-tune ABC and Treg homeostasis without triggering the severe developmental and immune dysregulation seen in complete knockout states. Future work should dissect how graded changes in PLCγ2 activity—rather than complete loss—affect ABC biology, Treg homeostasis, myelin maintenance, and microglial states over the lifespan and in disease models, and should test whether targeted modulation of specific PLCγ2-dependent pathways can harness the benefits suggested by human genetics without recapitulating the detrimental consequences observed in full Plcg2 deficiency.

## List of Abbreviations

PLCγ2: Phospholipase C gamma-2
PIP_2_: Phosphatidylinositol-4,5-bisphosphate
DAG: Diacylglycerol
KO: knockout
HET: heterozygous
WT: wild-type
APLAID: autoimmune PLCγ2-associated antibody deficiency and immune dysregulation
CVID: common variable immunodeficiency
IP_3_: inositol trisphosphate
PKC: protein kinase C
BCA: bicinchoninic acid
TCEP: tris(2-carboxyethyl)phosphine hydrochloride
PCA: principal component analysis
NeuN: neuronal nuclei
MAG: myelin associated glycoprotein
GFAP: glial fibrillary acidic protein
ROI: region of interest
DALs: differentially abundant lipids
PLS-DA: partial least squares-discriminant analysis
DEGs: differentially expressed genes
LRT: likelihood ratio test
MgND: neurodegenerative microglia
DAM: disease associated microglia
DAPs: differentially abundant proteins
KEGG: Kyoto Encyclopedia of Genes and Genomes
BCAAs: branched chain amino acids

## Declarations

### Ethics approval and consent to participate

For animal studies, experiments were performed in accordance with the guidelines approved by the Institutional Animal Care and Use Committee (IACUC) of University of Texas Health San Antonio (UT Health SA). Animals and other experimental units were assigned randomly to the experimental groups.

### Consent for publication

Not applicable

### Availability of data and materials

The vast majority of data generated or analyzed during this study are included in this published article and its supplementary information files. However, additional datasets used and/or analyzed during the current study are available from the corresponding authors.

### Competing Interest Statement

The authors declare that they have no competing interests.

## Funding

This work was supported by the Alzheimer’s Association [grant number AARG-21-846012 to SCH and JPP], the National Institutes of Health (NIH) [grant number K01AG066747 to SCH, Administrative Supplement 25S1 of parent grant AG013319 to JPP, RL5 scholar of P30AG044271 (JPP), PREP Scholar of R25GM130437 (JGR)], and UT Health SA institutional startup funds (JPP and SCH), a UT Health SA Department of Microbiology, Immunology & Molecular Genetics Pilot Grant (AVG, JPP and SCH).

### Author Contributions

Eduardo Gutierrez Kuri (Investigation, Software, Formal Analysis, Visualization, Writing – Original Draft, Writing – Review & Editing), Juliet Garcia Rogers (Investigation, Formal Analysis, Writing – Review & Editing), Sabrina Smith (Investigation), Gabriela Campos (Investigation), Henry Miller (Software, Formal Analysis, Visualization), Savannah Barannikov (Resources, Investigation), Hu Wang (Investigation), Sammy Pardo (Investigation), Xianlin Han (Resources, Supervision), Kevin F. Bieniek (Resources, Supervision), Susan T. Weintraub (Resources, Supervision, Writing – Review & Editing), Ann V. Griffith (Conceptualization, Investigation, Visualization, Supervision, Funding Acquisition, Writing – Review & Editing), Sarah C. Hopp (Conceptualization, Methodology, Investigation, Formal Analysis, Visualization, Supervision, Project administration, Funding Acquisition, Writing – Original Draft, Writing – Review & Editing), Juan Pablo Palavicini (Conceptualization, Methodology, Investigation, Formal Analysis, Visualization, Supervision, Project administration, Funding Acquisition, Writing – Original Draft, Writing – Review & Editing)

## Acknowledgments

Lipidomics analyses were performed at the UT Health San Antonio Barshop Institute Functional Lipidomics Core partially supported by NIH [AG013319 (San Antonio Nathan Shock Center of Excellence in the Basic Biology of Aging) and AG044271 (Claude D. Pepper Older Americans Independence Center)]. Nanostring analysis was performed at the South Texas Alzheimer’s Center [AG066546].

---

**Figure S1. Sex-stratified body weight, blood glucose, spleen weight, and brain weight in Plcg2-deficient mice, including an independent cohort.**(**A**) Body weight in male and female Plcg2 WT, Het KO, and Homo KO littermates at the 3-month endpoint, combining untreated animals (Figure 1) and an independent PBS-treated control cohort. (**B**) Non-fasted blood glucose at the 3-month endpoint in male and female Plcg2 littermates of each genotype from both cohorts. (**C**) Dry spleen weight at the 3-month endpoint in male and female Plcg2 littermates of each genotype from both cohorts. (**D**) Hemicerebrum dry weight at the 3-month endpoint in male and female Plcg2 littermates of each genotype from both cohorts. Data are presented as mean ± SEM; individual points represent individual animals (males, squares; females, circles). Two-way ANOVA with factors genotype and sex was followed by Tukey’s post hoc tests assessing simple genotype effects within each sex; p values (adjusted for multiple comparisons) < 0.1 are indicated in the graphs (A–D).

**Figure S2. Absolute spleen cellularity and myeloid/B-cell counts in Plcg2-deficient mice.**(**A**) Total live splenocyte counts in WT and Plcg2 Homo KO mice. (**B**) Absolute number of CD19⁻CD11b⁺ myeloid/innate-enriched cells per spleen. (**C**) Absolute number of CD19⁺CD11b⁻ B cells per spleen. Data correspond to the same animals shown in Figure 2 and are presented as mean ± SEM; individual points represent individual animals (males, squares; females, circles). Statistical significance was assessed using unpaired two-tailed t tests when data passed normality and homoscedasticity tests (A), Welch’s t test when F tests indicated unequal variances between groups (C), or Mann–Whitney tests when data did not meet normality assumptions (B); p values < 0.1 are indicated in the graphs.

**Figure S3. Extended analyses of mitochondrial lipid and amino-acid catabolism in Plcg2-deficient brains.** (**A**-**D**) DIA-MS proteomics was used to quantify additional mitochondrial matrix enzymes, including ALDH9A1, ACSS1, RIDA, and HMGCS1, in cerebrum from Plcg2 WT, Het KO, and Homo KO mice, with normalized protein abundances shown for each genotype. (**E**) Targeted LC/MS acylcarnitine metabolomics was used to assess correlations between individual acylcarnitine species and Plcg2 genotype; bar plots display correlation coefficients and corresponding p-values for selected short-, medium-, and long-chain acylcarnitines. Data are presented as bar plots with each point representing an individual animal; statistical models and multiple-testing corrections used for proteomic and metabolomic analyses are described in the Methods.

**Figure S4. Selective PIP_2_ depletion with preserved PIP_1_, PIP_3_, and DAG subclass levels in Plcg2-deficient cerebrum.** (**A**-**C**) Total levels of the abundant myelin-enriched species PIP_2_ 18:0–20:4, total PIP_1_, and total PIP_3_ were quantified by multidimensional shotgun lipidomics and expressed as nmol/mg protein. (**D**-**F**) To assess potential downstream effects on PLCγ2-derived diacylglycerol, total DAG was partitioned into saturated, monounsaturated, and polyunsaturated subclasses; none of these saturation-based DAG subclasses differed significantly by genotype. Bars show mean ± SEM; individual points represent individual animals (males, squares; females, circles). FDR-adjusted p values from linear models adjusted for sex are provided in the graphs.

## Notes

### Competing Interest Statement

The authors have declared no competing interest.

