## Supplementary figures and images for "PLCγ2 deficiency compromises systemic immune tolerance and erodes myelin homeostasis while enhancing oxidative metabolism in the mouse brain"

### Supplemental Figure 1

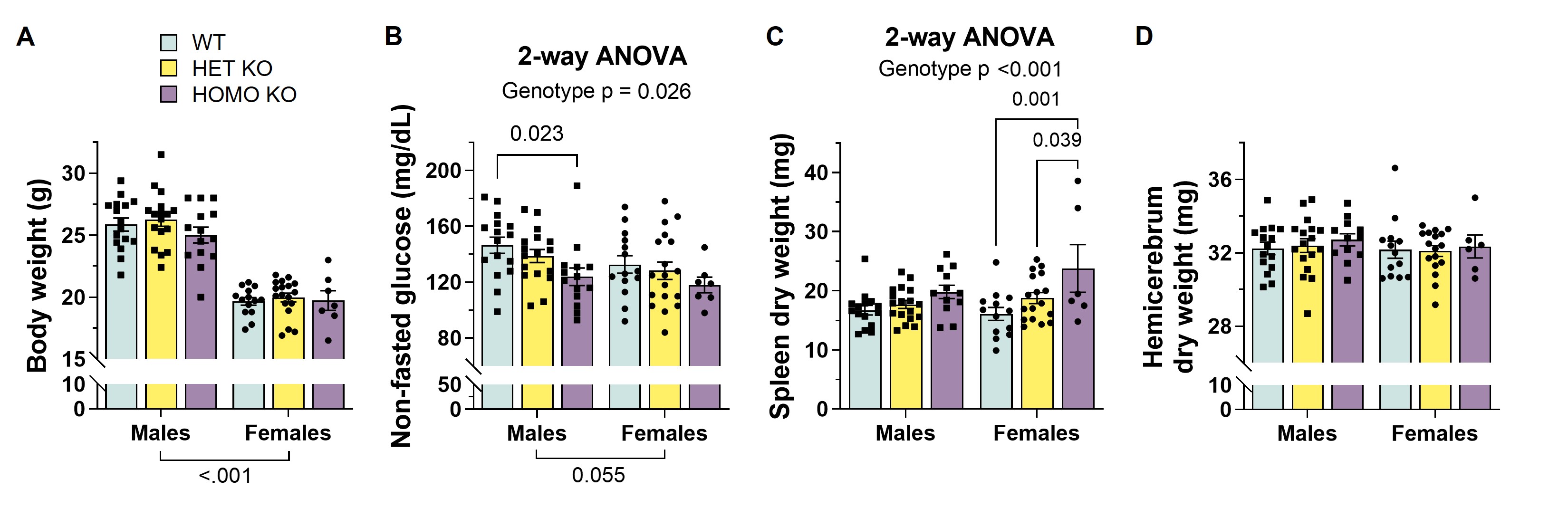

### Supplemental Figure 2

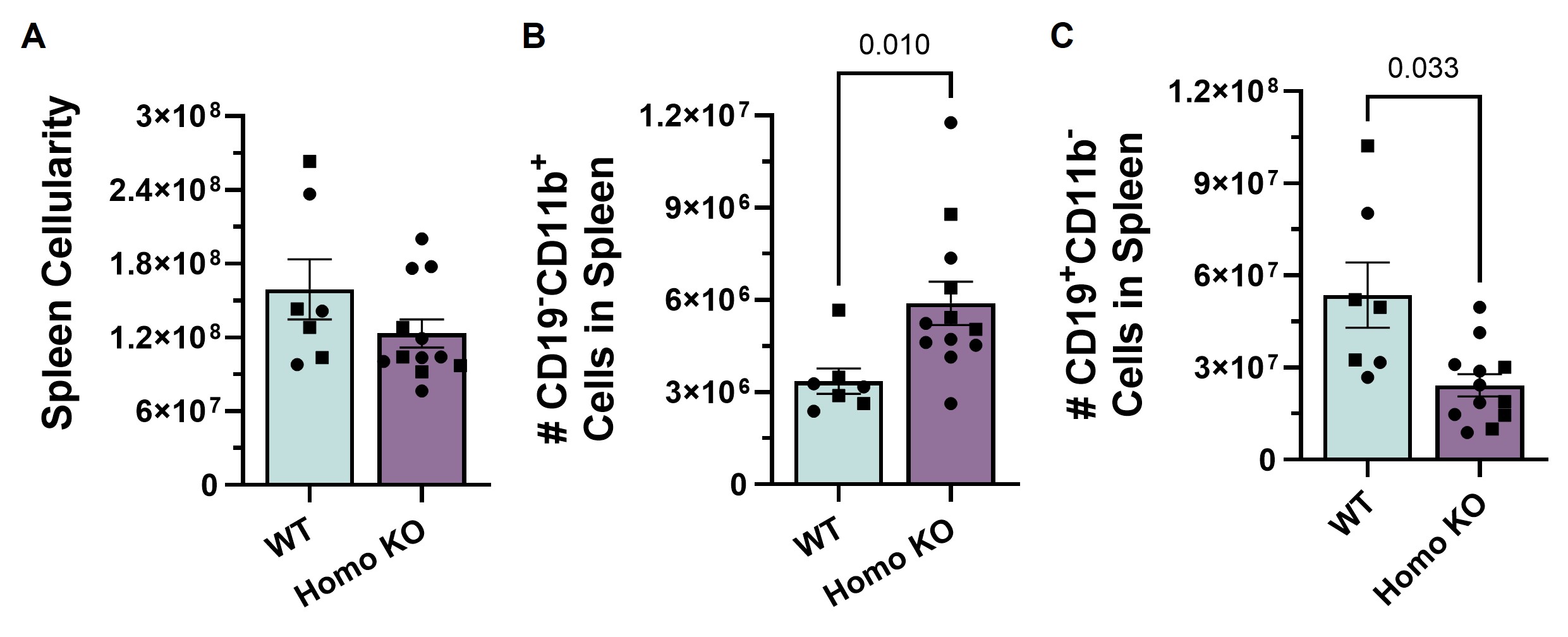

### Supplemental Figure 3

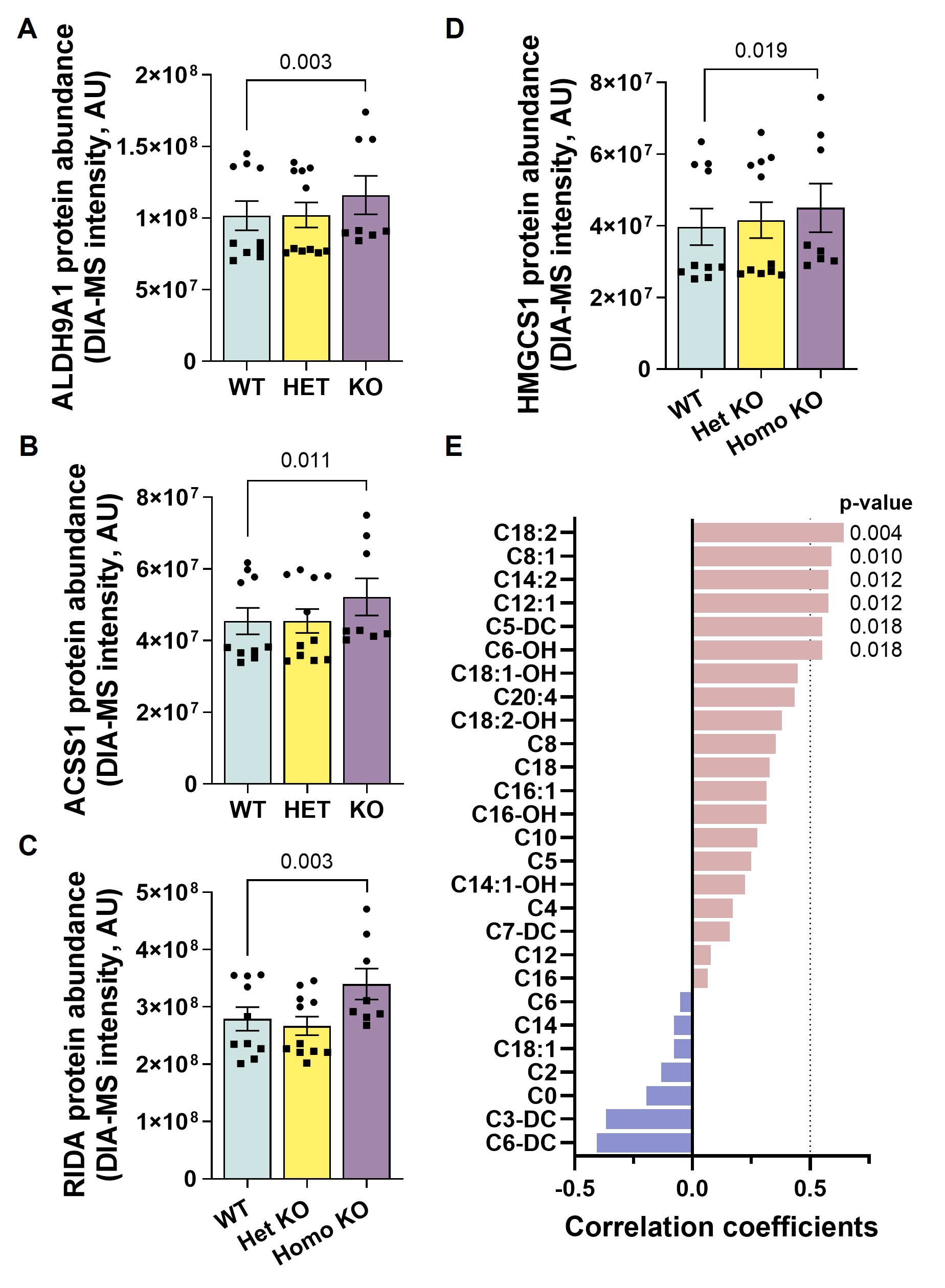

### Supplemental Figure 4

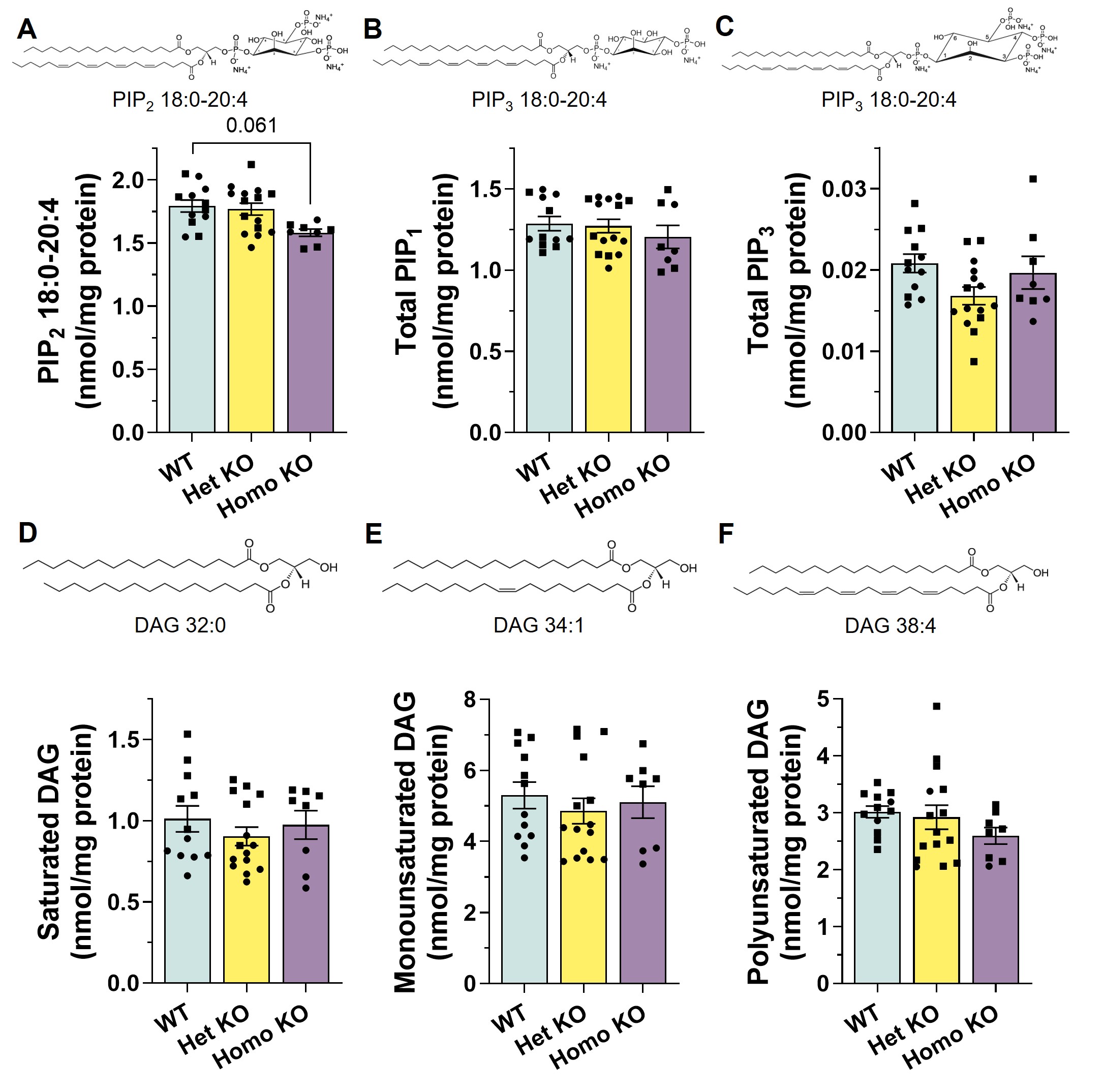

### Supplemental Table 1

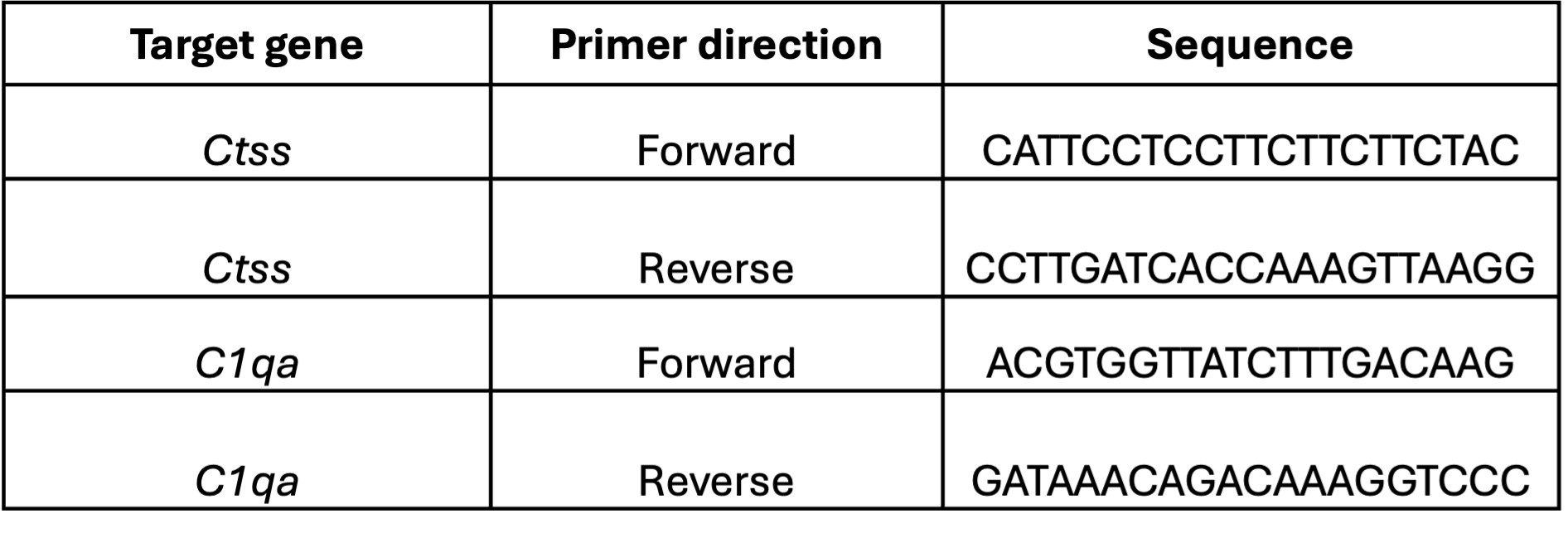
